# Definition and discovery of tandem SH3-binding motifs interacting with members of the p47^phox^-related protein family

**DOI:** 10.1101/2025.09.15.676255

**Authors:** Zsofia Etelka Kalman, Tamas Lazar, Laszlo Dobson, Rita Pancsa

## Abstract

SH3 domains are widespread protein modules that mostly bind to proline-rich short linear motifs (SLiMs). Most known SH3 domain-motif interactions and canonical or non-canonical recognition specificities are described for individual SH3 domains. Although cooperation and coordinated motif binding between tandem SH3 domains has already been described for members of the p47^phox^-related protein family, individual cases have never been collected and analyzed collectively, which precluded the definition of the binding preferences and targeted discovery of further instances. Here, we apply an integrative approach that includes data collection, curation, bioinformatics analyses and state-of-the-art structure prediction methods to fill these gaps. We define the optimal binding preference of tandemly arranged SH3 domains as [PAVIL]PPR[PR][^DE][^DE], and propose potential new instances of this SLiM among the family members and their binding partners. Structure predictions suggest the possibility of a novel, reverse binding mode for certain motif instances. A search of the human proteome with the sequence signatures of SH3 tandemization and follow-up structure analyses suggest that SH3 tandemization could be specific for this family. In all, our comprehensive analysis of this unique SH3 binding mode enabled description of the binding preference and identification of novel interesting cases proposed for experimental validation.

## Introduction

SH3 domains are small globular protein modules that are widespread interaction specialists. They represent one of the most abundant protein domain families: there are several hundred SH3 domain-containing proteins in the human proteome, many of them harboring more than one such domain [1]. They function in diverse signaling pathways, most often in membrane receptor-associated signaling pathways/networks. The majority of them are kinases or multi-domain adaptor, scaffold or effector proteins [1]. The most prominent function of SH3 domains is to specifically recognize (mainly proline-rich) peptide motifs in proteins, so-called short linear motifs (SLiMs), but through the years they turned out to be implicated in diverse recognition modes, including domain-domain, domain-RNA and domain-lipid interactions [1,2]. They are central players in the human protein-protein interaction (PPI) network, mediating the assembly of protein complexes through PPIs, as well as by enabling switch-like autoregulatory mechanisms through controllable intrachain domain-motif interactions [3–5]. Recently, several SH3-mediated multivalent PPIs were reported to drive liquid-liquid phase separation and thereby contribute to the formation of functionally specialized biomolecular condensates that confer efficiency and fidelity on cellular signaling [6–8].

Regarding their recognition specificities, SH3 domains are highly versatile [2,9]. A plethora of low-throughput and high-throughput (mainly phage display-based [10]) studies aimed at identifying the peptide-binding specificities of individual SH3 domains, leading to the description of canonical [11– 13] and non-canonical [10,14–16] binding specificities in both forward and reverse orientations [11,12]. The SH3-peptide complex structures helped the elucidation of some important specificity determinants of SH3 domains, i.e. the residues that comprise the shallow hydrophobic pockets of their binding cleft, which contribute to the binding of peptides that typically adopt a polyproline type II helix conformation [12,13]. Also, SH3 domain-mediated interactions turned out to be highly regulated by phosphorylations affecting the recognized motifs within binding partners or the peptide binding clefts of the SH3 domains themselves [17]. The SH3 domains of the human proteome were comprehensively compiled and classified according to different aspects, and their hitherto described binding specificities were also collected, summarized and analyzed [1,9]. Yet, this comprehensively compiled data clearly highlights that we are far from completely understanding the functioning of these multifaceted protein modules, and that the specificities and cellular binding partners of the majority of human SH3 domains remain undiscovered.

Besides falling short of the comprehensive description of binding specificities for all individual SH3 domains, another aspect of SH3 domain functioning that remains clearly understudied is a possible cooperation between different SH3 domains within the same protein or between different proteins. The paucity of such insights mainly stems from technical difficulties. In multi-domain modular proteins, globular domains are usually connected by disordered linker regions. While this arrangement allows for the free movement, rotations and independent functioning of neighboring domains, and therefore it is highly advantageous in cells, the increased length and conformational freedom of modular proteins make their *in vitro* experimental handling and characterization highly problematic. Due to this reason, the functioning/interactions of modular proteins are usually studied through separately investigating the binding properties of the individual protein modules. While this approach can certainly eliminate many technical difficulties and bring valuable insights, it is insufficient for uncovering functions/interactions that require cooperation between different modules of a protein.

The best-described case of cooperation between neighboring, tandemly arranged SH3 domains is that of p47^phox^ (gene name: NCF1). This protein regulates the assembly and activation of the phagocyte NADPH oxidase complex that catalyzes the conversion of oxygen to superoxide and other reactive oxygen species (ROS), and which is composed of membrane-embedded and cytoplasmic components [18]. The cytoplasmic p47^phox^ adaptor protein has an autoinhibited, closed conformation wherein its tandem SH3 domains (tSH3s) associate and form a common, composite binding groove that interacts with an autoinhibitory motif [18–20], a mode of binding that was also termed the superSH3 binding mode [21]. The composite ligand binding groove is formed by the conventional ligand binding grooves of the N- and C-SH3 domains, which bind the motif in opposite orientations (termed as plus/minus orientations) [20]. The core motif inserting into the composite binding groove adopts a polyproline type II (PPII) helix conformation, similarly to canonical SH3-binding motifs [19,20]. Several phosphorylations of serine residues in the vicinity of the autoregulatory motif are required to open up this closed conformation [22,23] and enable association of the tSH3s of p47^phox^ to a similar motif within p22phox (gene name: CYBA) [18,19,24], thereby forming a bridge between the membrane-embedded and cytoplasmic components of the NADPH oxidase and activating the complex [18,19].

The p47^phox^-related organizer superfamily of proteins has 5 members, p47^phox^ (gene:NCF1), p41^phox^ (gene:NOXO1), p40^phox^ (gene:NCF4), TKS4 (gene:SH3PXD2B) and TKS5 (gene:SH3PXD2A), with all of them harboring an N-terminal PX domain and one or more SH3 domains. For simplicity, we will refer to the proteins by their gene names from now on. SH3PXD2B and SH3PXD2A will be referred to by their alternative gene names: TKS4 and TKS5, and the protein family will be referred to as the NCF1 family. Potential association and motif-binding of tandem SH3 domains has only been scarcely studied in other members of the superfamily [25–27] and, to our knowledge, it has never been investigated in proteins outside of this superfamily, although there are many proteins harboring multiple consecutive SH3 domains. Studying this binding mode experimentally is highly challenging, as the association of the tSH3s is very weak: even for NCF1, the two domains were only able to associate when linked (ensuring high local concentration) but not when separately placed into the solution [19]. Also, binding of the peptide motif has a clear stabilization effect on the complex [19], therefore studying a construct with a pair of linked SH3 domains alone might still not be sufficient to detect the association.

In this study, we first comprehensively analyze available literature evidence on the association and motif binding of tandem SH3 domains in members of the NCF1 family to then use the accumulated motif instances, their structural properties (if available), evolutionary conservation patterns and mutation data to define the underlying binding determinants as a novel short linear motif (SLiM). In this step, we follow the strategy that mirrors the annotation procedure for new motif classes in the Eukaryotic Linear Motif (ELM) database [28,29]. The resulting motif definition is then used to propose new motif instances in the family members and their binding partners. Furthermore, we aim to establish the sequence requirements (on the domain side) for the association of tandemly arranged SH3 domains in members of the NCF1 family using the available complex structures, as well as conservation patterns within the family and through evolution. Using these signatures, we screen the human proteome to see if we can identify other multi-SH3 domain proteins whose tandemly arranged SH3 domains could potentially associate and act in cooperation for motif binding.

## Results

### Multiple lines of evidence support the cooperation of tandem SH3 domains in motif binding for members of the p47^phox^-related organizer superfamily

Among the five members of the p47^phox^-related organizer superfamily (referred to as the NCF1 family), NCF4 is the only one which has a single SH3 domain. Other members have multiple SH3 domains, of which the first two are in proximity in sequence, connected only by a relatively short linker (**Figure 1**). In the following sections, we discuss the literature evidence supporting the cooperation of tandem SH3 domains (tSH3s) in motif binding for the four family members and collect all instances of the tSH3-binding motifs hitherto described (**Table 1**).

**Table 1:**
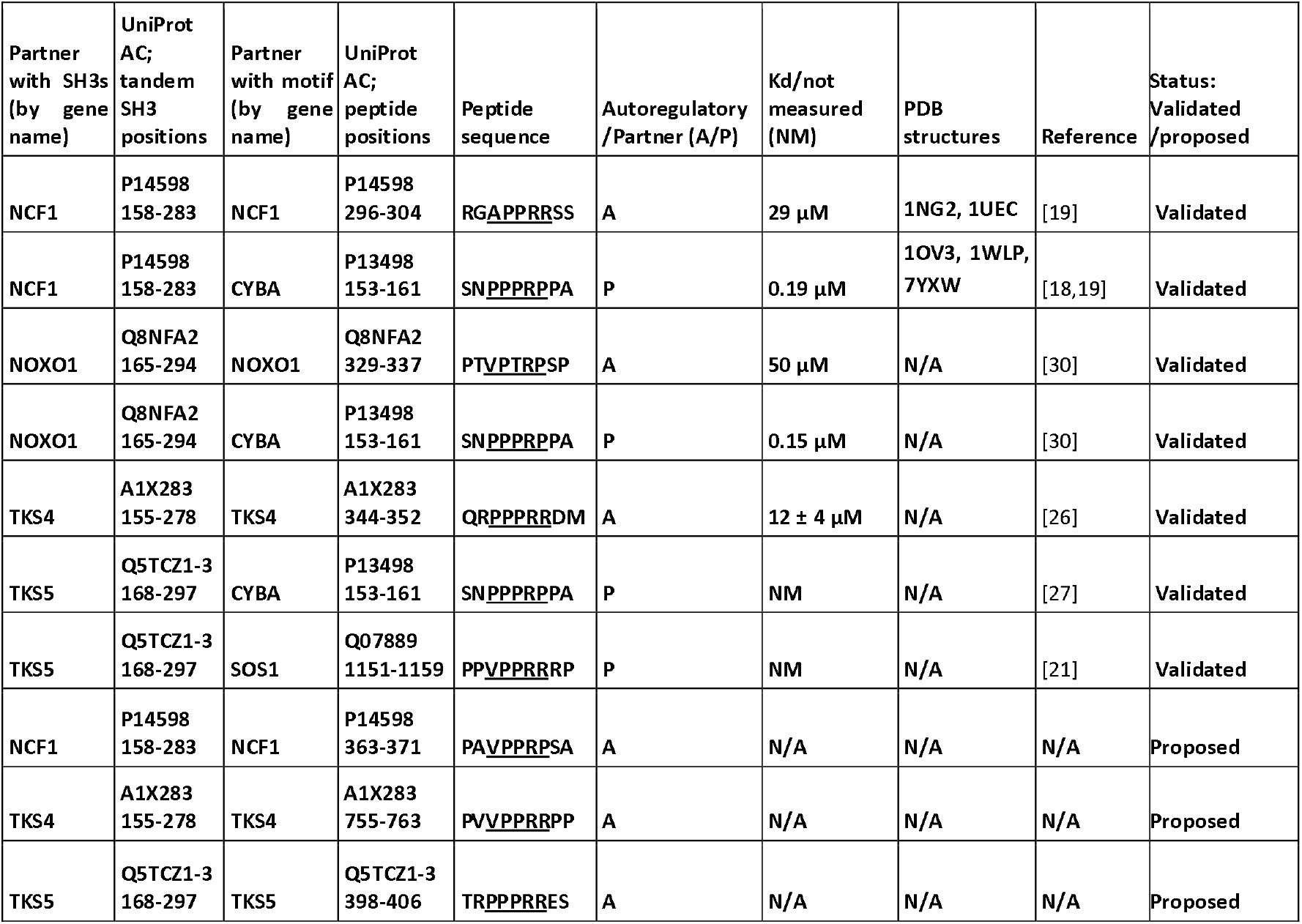
Experimentally validated and proposed tandem SH3-binding motifs of the NCF1 family.

**Figure 1:**
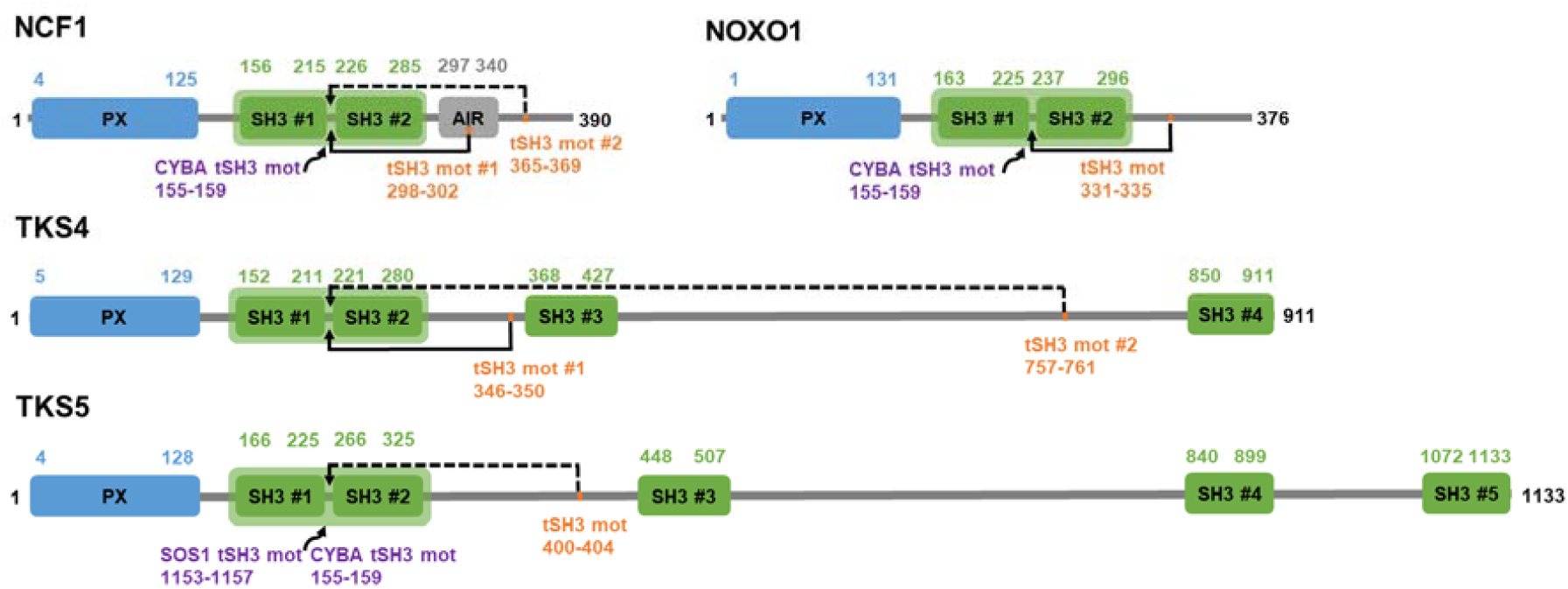
Domain maps of the NCF1 family members with more than one SH3 domain. Domain maps of NCF1, NOXO1, TKS4 and TKS5. PX domains are depicted in blue, SH3 domains in green. Tandemized SH3 domains are marked with a light green background. Autoregulatory tandem SH3-binding motifs (tSH3 mot) are marked with orange, among these, the experimentally verified cases are connected to the tandem domains by continuous lines, while the proposed motif instances are connected by dashed lines. The gene names of partner proteins that are experimentally proven to bind to the tandem SH3s, along with the positions of their tSH3-binding motifs are indicated in purple.

NCF1, the eponym of the family, is the major cytoplasmic regulatory subunit of the NADPH oxidase complex [18]. It has an autoinhibited closed conformation where the tSH3s form a composite binding cleft to bind to a C-terminal proline-rich “APPRR” motif within the autoinhibitory region (AIR) [19]. This conformation can be resolved by phosphorylations directly downstream of the motif [22], leading to an active, open conformation, where the tSH3s of NCF1 can bind to a similar “PPPRP” motif within one of the catalytic subunits of the oxidase, namely p22^phox^ (gene: CYBA), which activates the oxidase complex [18,19,24]. The two motifs are very similar, and both SH3 domains of NCF1 are required for the binding in both cases [19].

NOXO1 is a homologue of NCF1 with 22% identity that functions similarly in the activation of superoxide-producing NADPH oxidases. Since it lacks the autoinhibitory region (AIR) present in NCF1, it was first considered as a constitutively active organizer of the NADPH oxidase [18,25]. In contrast to full-length NCF1 that is in an autoinhibited form, full-length NOXO1 is able to interact with CYBA through its tSH3 domains. The NOXO1-CYBA interaction is similarly disrupted by the chronic granulomatous disease (CGD)-associated P156Q mutation in the CYBA motif 155-PPPRP-159, as the NCF1-CYBA interaction [18,25]. However, later on a relatively weak autoinhibitory interaction was uncovered between the tSH3s of NOXO1 and its C-terminal Pro-rich region, namely residues 325-TAPPPTVPTRPS-336 [30,31]. Albeit weak, this autoinhibitory interaction in NOXO1 was shown to weaken the binding to CYBA. Pro to Ala mutation of three residues (P329, P332 and P335) abrogated autoinhibition, leading to CYBA binding with higher affinity [30]. It is important to note that the identified segment harbors the sequence “VPTRP” that is very similar to the motif bound within CYBA and to those bound by the tSH3s of other NCF1 family members (see below). Interestingly, the same Pro-rich region of NOXO1 is known to mediate the interaction with the SH3 domain of NCF2/NOXA1 [25].

TKS4 and TKS5 represent a distinctly different subfamily within the p47^phox^-related organizer superfamily. They are large, modular adaptor proteins that contribute to the formation of podosomes and invadopodia and thus are implicated in cell motility, including the migration of cancer cells during metastasis [32–34]. Interestingly, TKS5 has been demonstrated to fulfil a similar role to NCF1 by binding to CYBA and thereby facilitating ROS production during invadopodia formation [27].

The N-terminal part of TKS4 closely resembles NCF1 and was demonstrated to mediate similar intramolecular interactions as shown in a study by Merő B. and colleagues [26]. Their *in vitro* binding assays and SAX results support that TKS4 has an autoinhibited conformation wherein the 3rd SH3 domain binds to the N-terminal PX domain and the tandemly arranged 1st and 2nd SH3 domains engage with a highly conserved “PPPRR” motif located between residues 346 and 350 [26]. They explicitly show that the individual SH3 domains are incapable of binding to this region, so the tandem arrangement is strictly required. Additionally, a phosphomimetic mutation was introduced into the peptide sequence near the identified motif, but it did not considerably affect binding affinity, implying that the regulation may differ from that in NCF1. The authors only used constructs covering the first part of the protein, regions downstream of the 3rd SH3 domain have not been investigated [26].

The tandem SH3 binding mode has also been suggested for an isoform of TKS5 wherein the linker connecting the first two SH3 domains is shorter than in the canonical isoform (see UniProt: Q5TCZ1-3) [21]. Also, TKS5 was reported to interact with the same region of CYBA as NCF1 and NOXO1 (155-PPPRP-159), moreover, mutational analyses indicated that the first two (the tandem) SH3 domains are both implicated in binding and the binding is compromised by the same disease-associated mutation (P156Q mutation in CYBA) that has also been reported to affect the binding of NCF1 [27] and NOXO1 [25]. Furthermore, Rufer AC *et al*. demonstrated some isoform-specific protein-protein interactions for TKS5. In their peptide spot membrane assay, four peptides of SOS1 were exclusively bound by the tSH3s of the short TKS5 isoform, resulting in a strong signal (peptides 2, 6, 14 and 25) [21]. Peptide 14 covering human SOS1 residues 1145-1164 contains “VPPRR”, so we also considered it as a validated instance of the PPRR motif (Table 1). We cannot explain the exclusive binding of peptides 2, 6 and 25 to the tSH3s of the short TKS5 isoform, because those do not contain anything resembling a PPRR motif. In the same study, DNM1 and DNM2 peptides were also tested similarly. Only 1 DNM1 and 2 DNM2 peptides produced a weak signal specific for binding the tSH3s. However, based on affinity data obtained for individual and tSH3s in follow-up ITC titration experiments, the authors argued against the synergistic binding mode in the case of the TKS5-DNM1/2 interactions [21].

### The binding preference of the tSH3s of the NCF1 family: definition of the tandem SH3-binding motif

By taking into account the contacts in the available complex structures, constraints imposed by the PPII helix conformation on the motif residues [35], binding strengths, evolutionary conservation signatures, data from mutational analyses, and experimentally verified disease mutations, we here make an attempt to define the crucial binding determinants of tandem SH3 domains (*See Graphical abstract*). We describe the motif as a regular expression that is commonly used for defining short linear motifs (SLiMs) [28,29] and can be used for identifying novel motif instances in protein sequences.

Based on the vertebrate alignments of known motif instances (**Figure S1**) and DSSP secondary structure annotations in the available X-ray structures, we found that five consecutive residues show high conservation and a tendency to form PPII helix conformation, thereby forming the core of the proposed motif (**Figure 2**). A strong/strict version of the motif core can be defined as [PAVIL]PPR[PR]. In the available X-ray structures depicting the interaction of the NCF1 tSH3s with the autoregulatory motif (PDB IDs:1NG2 and 1UEC) or the motif in the partner CYBA (PDB IDs:1OVC3 and PDB:7YXW), the motif core forms a polyproline type II helix (PPII) structure [20] with minor variations in the starting and finishing positions, as detected by DSSP [36]. In the first position of the core motif, the selection of residues (Pro and hydrophobic residues) was defined based on the vertebrate alignments of the validated motif instances (**Figure S1**) and the fact that this position is already part of the PPII helix observed in the structures. Here, Pro, Ala, and Val are seen most frequently, while the larger hydrophobic residues, Leu and Ile, are observed less frequently. In the second position, Pro seems to be strictly required based on several observations: 1) there is absolutely no variation in the alignments of the motifs in this position, 2) the chronic granulomatous disease-associated mutation within CYBA (P156Q) hits this second position and is known to abrogate binding, 3) the Pro in this position (equivalent to P299 within the AIR of NCF1) makes contacts with both domains of the tandem SH3 [19,20]. In the third position, Pro seems to be strongly preferred for strong binding, based on its invariant conservation in the strong motif instances (NCF1 AIR, CYBA, TKS4 autoregulatory motif, SOS1 motif). Also, this Pro (equivalent to P300 within the AIR of NCF1) makes several contacts with a hydrophobic pocket of the first SH3 domain of tSH3s [19,20]. In the fourth position, Arg is invariably conserved among vertebrates for all the motif instances, and it is known to make both hydrophobic and electrostatic contacts with the second SH3 domain [19,20], so it seems to be strictly required for binding. In the fifth position, either Pro or Arg seems to be allowed, since only these two residues can be seen in the motif alignments. The residue in this position contacts the first SH3 domain and it was predicted as the last residue of the PPII helix in one of the available structures of the complex (PDB ID: 7YXW). Finally, some restrictions can be defined for the C-terminal flanking residues of the motif. In the first and second positions immediately following the motif core (6th and 7th positions) negatively charged residues are not allowed because they definitely weaken the interaction. This is evident because phosphorylation or a change to phosphomimetics [22] of the serine residues in these positions of the NCF1 autoregulatory motif was demonstrated to weaken the binding [19]. Also, a negatively charged residue at position 6 would induce charge repulsion with a negatively charged residue of the 2nd SH3 domain (Glu241 in NCF1) [19]. Based on these considerations, a strong tandem SH3-binding motif can be defined as [PAVIL]PPR[PR][^DE][^DE]. In AlphaFold (AF)-predicted tandem SH3-motif complexes of all the known or proposed motif instances in **Table 1** that adhere to this strong motif definition, the motifs are perfectly aligned (*see Materials and methods for details on the structure predictions;* **Figure 2A**). We use the predicted complexes for consistency, but they correspond precisely to the respective PDB structures, where available (**Figure S2**).

**Figure 2:**
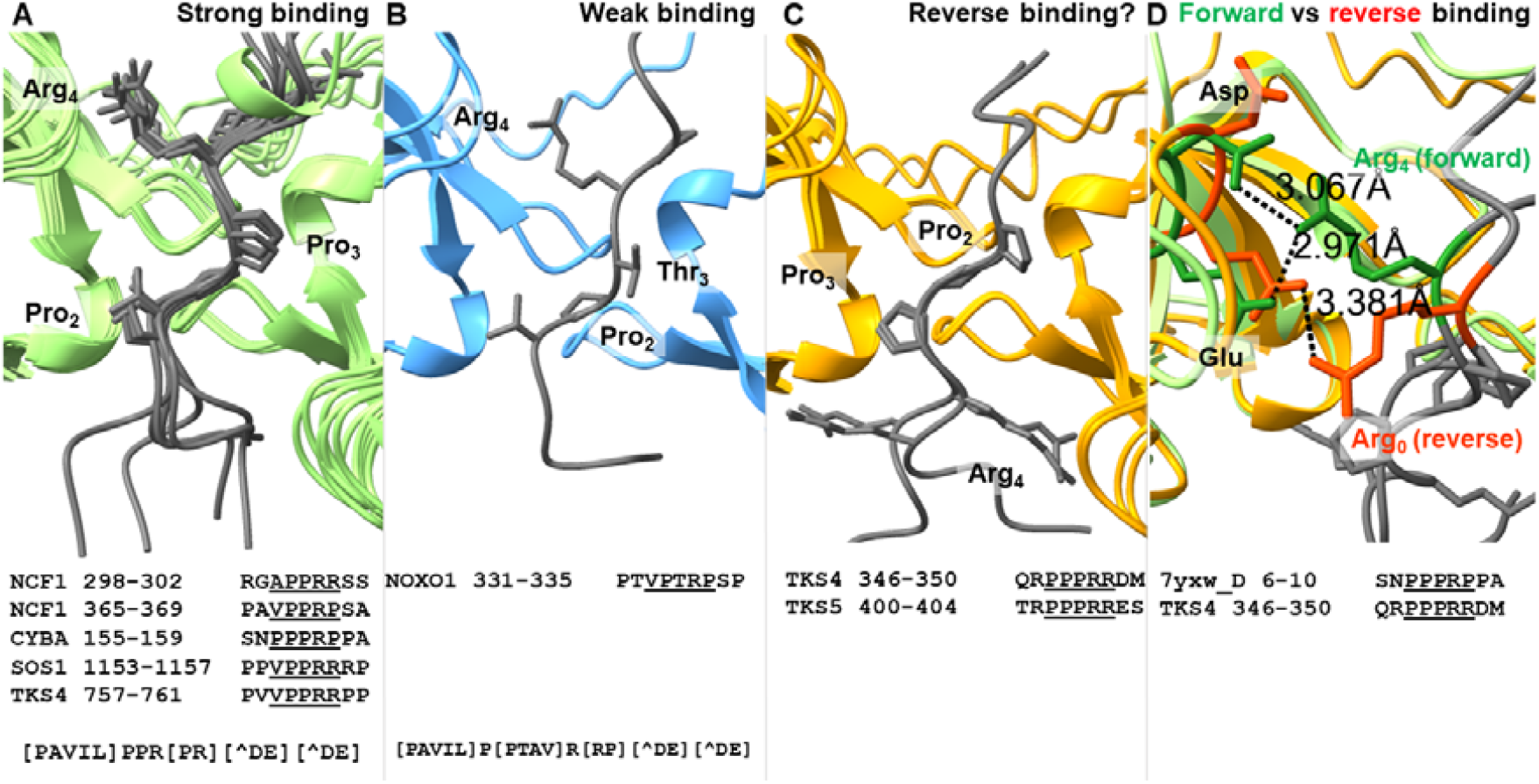
Different binding modes of the tandem SH3-binding motifs. AlphaFold-predicted complex structures suggest different binding modes based on motif sequence characteristics. A) All motifs fitting the strong motif definition bind very similarly (this is also evident based on the experimentally determined structures listed in **Table 1**). B) The weak binder NOXO1 C-terminal motif that has a Thr in the 3rd position of its core binds in an almost identical manner as the strong binders, which supports the validity of the weak motif definition. C) A reverse binding mode is predicted for a pair of homologous motifs in TKS4 and TKS5, where a negatively charged residue follows the core motif, but an Arg occupies position 0, directly preceding the core motif. D) Comparison of the positioning of Arg4 in the forward binding mode versus Arg0 in the reverse binding mode and their distances to the Glu and Asp residues of the C-double negative signature (defined in a following section) on the domain side.

Some validated motifs do not adhere to this strong motif definition (**Table 1**). The weakly binding NOXO1 C-terminal motif contains a Thr at the 3rd motif position, and the vertebrate alignment of this motif suggests that Ala and Val can also be accommodated here (**Figure S1**). Importantly, these residues are compatible with the residues observed in middle or penultimate positions of PPII helix structures [35] (in the two tandem SH3-binding motifs with available structures (**Table 1**), the detected PPII helix extends up until the 4th or 5th position as judged by DSSP [36]). Furthermore, based on AF predictions, this weaker motif can bind to the tSH3s of NOXO1 in a conformation that is very similar to how the strong motifs bind to their respective tandem SH3s (**Figure 2B**). Therefore, a less strict version of the motif can be defined as [PAVIL]P[PTAV]R[RP][^DE][^DE].

In the validated TKS4 autoregulatory motif (residues 346-350), as well as in the homologous motif of the closely related TKS5, there is a conserved negatively charged residue in the 6th position, the position directly following the core motif (**Table 1, Figure S1**), and therefore these motifs do not fit the strong motif definition defined above. A negatively charged residue in this position was already proposed to weaken the interaction based on several pieces of evidence [19]. The interaction of the TKS4 peptide was confirmed by biophysical experiments, suggesting a relatively weak affinity (Kd = 12 μM). Interestingly, AF2 predicts a reverse binding mode for these two similar motifs in all the five models of the output (**Figure 2C**). Out of curiosity, AF3 was also used on the autoregulatory cases, and the resulting complexes support the forward binding mode for motifs fitting the strong motif definition and the reverse binding mode for the TKS4 and TKS5 autoregulatory motifs that reside ahead of their 3rd SH3 domains (**Figure S3**). The validity of these predictions cannot be judged based on the currently available data, since the experiments performed by Merő *et al*. do not inform on the directionality of binding [26] and structural evidence is not available. For these two peptides, the reverse binding mode could be enabled by Arg residues occupying the 0 position (the one directly preceding the core motif). In the predicted reverse structure, these arginines occupy a similar position as the strictly required Arg in position 4 of the forward-binding strong motifs (**Figure 2D**). Still, while the latter can form salt bridges with both of the adjacent negatively charged residues within the 2nd SH3 domain of the tandem (alignment positions 267 and 268 in **Figure 3** below), Arg0 of the predicted reverse motif is a bit shifted and therefore can only contact the second acidic residue of the two (**Figure 2D**). TKS4 and TKS5 are closely related, and the region harboring this potential autoregulatory motif is fully conserved in their vertebrate alignments (the conservation pattern is not island-like as usually seen for functional SLiMs [37], and there is no sequence variation in the associated positions). Therefore, a more general reverse tSH3-binding motif cannot be defined based on these two cases.

**Figure 3:**
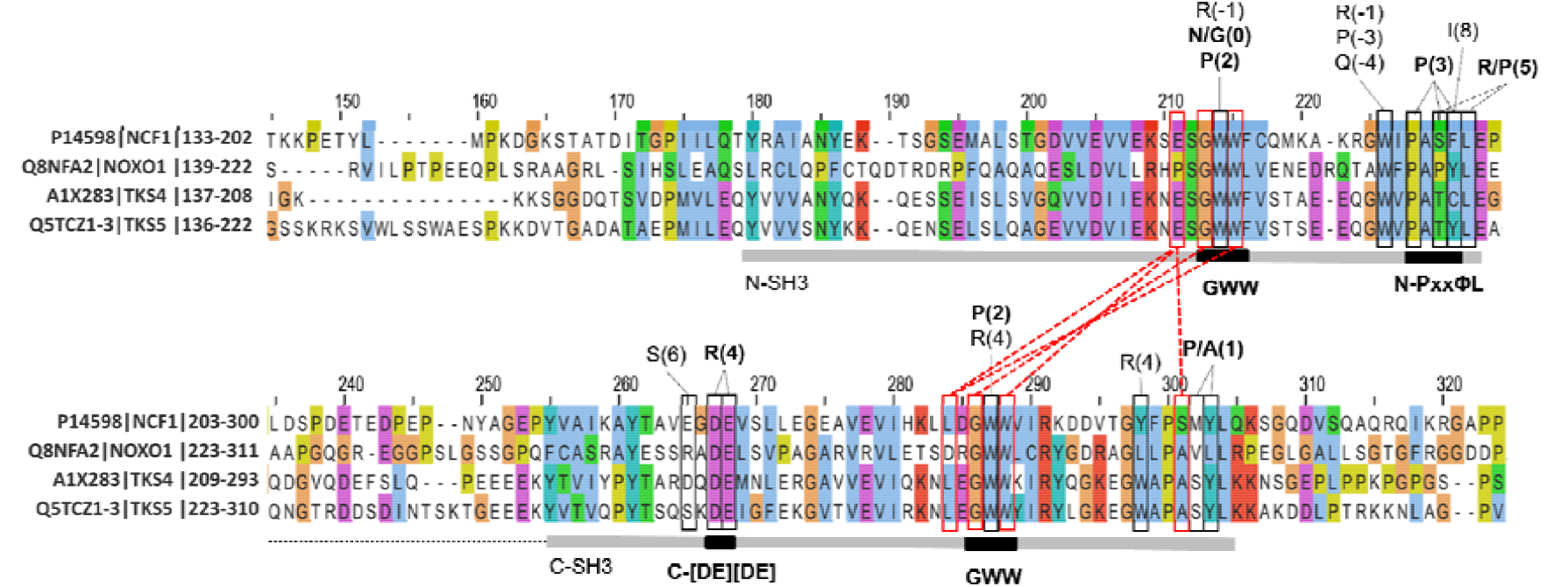
Sequence signatures of tandemization and motif binding within the tSH3s of the four family members. Positions suggested to be important in tandemization are marked with red boxes (based on [19,20]). Amino acids that participate in binding the partner peptides within the available complex structures (PDB IDs: 1NG2, 1UEC, 1OV3, 1WLP, 7YXW) are marked by black boxes. The residues of the contacting peptides are marked above the contacted alignment positions (bold letter: found in three or more structures, normal: found in at least two structures). The numbering of peptide/motif residues follows the one introduced in the previous section on the definition of the tandem SH3-binding motif, so, for instance, R(4) stands for the conserved Arg in the 4th position of the motif. The domain and linker boundaries, as well as the sequence signatures of tandemization and cooperative motif binding, are indicated in gray and black below the alignment.

### Detection of tandem SH3-binding motif candidates within the NCF1 family and their binding partners

We propose three potential novel autoregulatory motif candidates residing in intrinsically disordered regions (IDRs) [38,39] of NCF1 family proteins (see the last three lines of **Table 1**), of which two fit the strong motif definition proposed above, and one is a candidate for the reverse binding mode. A conserved, C-terminal, proline-rich motif of NCF1 “365-VPPRP-369” (**Table 1**, line 8) shows homology to the validated C-terminal autoregulatory motif of NOXO1, moreover it is likely a stronger binder since it has an optimal Pro residue in the third motif position. We are not aware of an alternative intramolecular interaction within NCF1 wherein its tSH3s interact with this conserved Pro-rich region instead of the well-studied AIR motif. However, we propose that such an interaction might exist because the “VPPRP” motif is highly similar to the well-studied AIR and CYBA motifs, and contains all the key residues known to be sufficient for binding to the tSH3s of NCF1 [19]. Furthermore, these C-terminal motifs in NOXO1 and NCF1 both contain a Ser in the position directly following the core motif, which suggests that they could also be regulated by phosphorylation (as the AIR-resident motif). Accordingly, the respective Ser residue can be phosphorylated in both proteins based on PhosPhoSite+ annotations [40].

Additionally, we propose that TKS4 might have a 2nd, C-terminal autoregulatory motif (**Table 1**, line 9). Notably, in the AlphaFold2 [41,42] structure of TKS4, which became available since the publication by Merő *et al*., the tSH3s bind to a “757-VPPRR-761” motif that differs from the one proposed by the in vitro experiments [26]. This alternative autoregulatory motif fits the strong motif definition, and accordingly, it is predicted to bind in the forward mode (and likely with high affinity), in contrast to the experimentally studied TKS4 autoregulatory motif, which is predicted to bind in reverse mode. Also, the proposed motif is located within the long IDR connecting the 3rd and 4th SH3 domains of TKS4. This region downstream of the 3rd SH3 domain of the protein was not covered by the *in vitro* studies, which could be the main reason why the interaction remained undiscovered.

To our knowledge, an autoregulatory intramolecular interaction has never been proposed for TKS5. Yet, a conserved “400-PPPRR-404” motif, located upstream of the 3rd SH3 domain (see **Figure 1** and **Table 1**, line 10) could likely serve as the autoregulatory module of the short TKS5 isoform (UniProt AC: Q5TCZ1-3), the one compatible with SH3 tandemization. At least one autoregulatory motif was described in all other NCF1 family members, so autoregulation is likely a conserved feature within the family. The functional importance of this motif is further supported by the fact that it is homologous to the validated autoregulatory motif of the closely related TKS4 [26]. Additionally, since the core motif is directly followed by an acidic residue, and preceded by an Arg, this motif is predicted to bind in the reverse binding mode, similarly to its TKS4 homolog (**Figure 2CD**).

To identify motif candidates within the known binding partners of the NCF1 family members, the strong motif definition ([PAVIL]PPR[PR][^DE][^DE]) was first searched in the whole canonical human proteome **(Table S1)** [43] and then the resulting 324 proteins were checked for an overlap with known binding partners of the NCF family members cataloged in BioGRID [44] and IntAct [45]. Predicted structural disorder [46] and sequence conservation were calculated to help evaluate the possible functionality of the identified hits [39]. The localizations of the proteins were also considered, since NCF1 family members are all cytoplasmic proteins that can be attached to the plasma membrane by their PX domains under certain circumstances. Besides self-interactions and partners already covered in **Table 1**, some interesting motif candidates could be identified (**Table 2, Figure S4**).

**Table 2:**
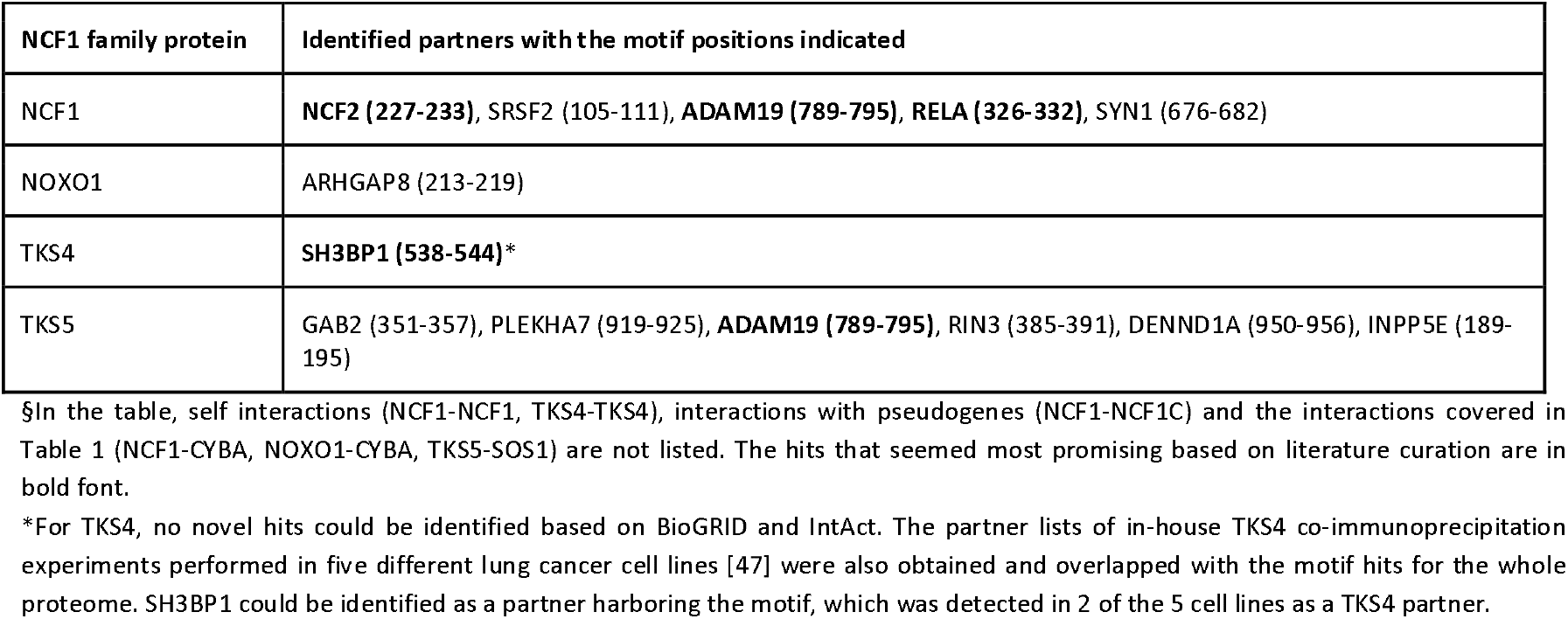
Partners of the NCF1 family proteins which contain at least one strong tandem SH3-binding motif.

| NCF1 family protein | Identified partners with the motif positions indicated |
| --- | --- |
| NCF1 | **NCF2 (227-233)**, SRSF2 (105-111), **ADAM19 (789-795)**, **RELA (326-332)**, SYN1 (676-682) |
| NOXO1 | ARHGAP8 (213-219) |
| TKS4 | **SH3BP1 (538-544)\*** |
| TKS5 | GAB2 (351-357), PLEKHA7 (919-925), **ADAM19 (789-795)**, RIN3 (385-391), DENND1A (950-956), INPP5E (189-195) |

§In the table, self interactions (NCF1-NCF1, TKS4-TKS4), interactions with pseudogenes (NCF1-NCF1C) and the interactions covered in Table 1 (NCF1-CYBA, NOXO1-CYBA, TKS5-SOS1) are not listed. The hits that seemed most promising based on literature curation are in bold font.
\*For TKS4, no novel hits could be identified based on BioGRID and IntAct. The partner lists of in-house TKS4 co-immunoprecipitation experiments performed in five different lung cancer cell lines [47] were also obtained and overlapped with the motif hits for the whole proteome. SH3BP1 could be identified as a partner harboring the motif, which was detected in 2 of the 5 cell lines as a TKS4 partner.

NCF2/p67^phox^ is a well-known interaction partner of NCF1 in the assembly of the NADPH oxidase, wherein the C-terminal SH3 domain of NCF2 is known to interact with a C-terminal Pro-rich motif in NCF1 [48], the one that we propose as a potential secondary autoregulatory site within NCF1 (**Table 1**, line 8). Since NCF2 has a Pro-rich disordered region with “PPPRPKT” between residues 227-233 that fits our strong motif definition, there could be a secondary interaction between the two proteins, where the tSH3s of NCF1 (after being released from autoinhibition) interact not only with the motif within CYBA [18], but also with a motif within the other cytoplasmic oxidase subunit, NCF2. Although an interaction between NCF1 SH3 domain(s) and NCF2 has been suggested [18], to our knowledge the mechanism and the binding site within NCF2 haven’t been precisely described yet. Based on our motif definition, a tandem SH3-binding motif maps to residues 227-PPPRPKT-233 of NCF2 and the key residues are almost fully conserved in vertebrates (**Figure S4**).

The interaction between NCF1 and RelA protein has been demonstrated using a range of interaction detection methods and is proposed to be important in the activation of the NF-κB pathway by IL-1 in endothelial cells [49]. The study demonstrated that the tandem SH3 domains of NCF1 are required for the interaction, and it narrowed down the interacting region within RelA to a larger Pro-rich IDR between the Rel homology and transactivation domains (region 301-431 in UniProt AC: Q04206) [49]. Based on our motif definition, the tandem SH3-binding motif can be precisely located to residues 326-PPPRRIA-332 of RelA. The key residues of the motif are generally conserved among placental mammals, only some Pro to Thr (or Ala) changes can be seen in the 3rd motif position (**Figure S4**), a change that could still be compatible with binding according to our weak motif definition.

The disordered cytoplasmic tails of ADAM family metalloproteinases were tested for interactions with a number of individual SH3 domains [50]. The 1st SH3 domain of NCF1 [50] and the 1st and 5th SH3 domains of TKS5 were found to bind to Pro-rich regions within ADAM19 [50,51] has a promising tSH3-binding motif between residues 789-PPPRPPP-795 (that is fully conserved among mammals (**Figure S4**)), so we propose that this potential interaction should be tested using tandemly arranged SH3 domain constructs.

SYN1 and SRSF2 do not seem to be real interaction partners of NCF1. For SYN1 only very weak binding was seen *in vitro* with the 2nd SH3 domain of NCF1 [52], with no biological relevance suggested for the interaction. The SRSF2 splicing factor is a primarily nuclear protein (in contrast to NCF1 which is cytoplasmic), and the NCF1-SRSF2 interaction was only detected in a single clone during a yeast two-hybrid screen [53]. The NOXO1-ARHGAP8 interaction is also not very well-supported, there is only one high-throughput affinity capture-MS analysis where this interaction was detected [54], but due to the characteristics of the methodology, the interaction could also be indirect.

SH3 domain-binding protein 1 (SH3BP1) is a GTPase-activating protein (GAP) inactivating the RAC1 GTPase at the leading edge of migrating cells [55], thereby facilitating the reorganization of the cytoskeleton at cell protrusions, such as podosomes, invadopodia and others. Its interaction with TKS4 has been detected by TKS4 co-immunoprecipitation experiments in different cell types [47]. The interaction is further supported by the fact that SH3BP1 and TKS4 share many common binding partners, including the adaptor proteins CIN85 and CD2AP [56,57], the SRC kinase, and the actin capping protein [47,57]. Therefore, cooperation between the two proteins in actin remodeling within podosomes and invadopodia is highly likely, and the detected tSH3-binding motif suggests a possible binding mechanism. However, the conservation of the potential tSH3-binding motif is not very strong (**Figure S4**), and the long, C-terminal IDR of SH3BP1 is so rich in prolines that there are plenty of potential SH3-binding sites in it (this is where its name “SH3 domain-binding protein 1” comes from). Thus, it could also be bound by any of the individual SH3 domains of TKS4, the binding preferences of which are poorly characterized [10].

GAB2, PLEKHA7, DENND1A and INPP5E have been suggested as TKS5 partners by a single study, where the interactome of cilia was characterized by high-throughput affinity capture-MS [58]. Since, to our knowledge, no other study suggested ciliary localization for TKS5, we are not sure about the validity of these interactions. Also, affinity capture-MS tends to detect indirect interactions. The detected potential tSH3-binding motif within GAB2 resides within a linear motif “PxRPPK” where GAB2 is known to be bound by the C-terminal SH3 domain of GRB2 [59]. Therefore, the evolutionary conservation of the potential tSH3-binding motif cannot be independently evaluated. The interaction between RIN3 and TKS5 has also been suggested based on a single high-throughput affinity capture-MS study [60]. The detected potential tSH3-binding motif within RIN3 overlaps with a validated SH3-binding site of CD2AP [61]. The interaction between RIN3 and CD2AP is well-characterized [60,61] and explains evolutionary conservation of the motif positions, therefore, again, validity of the potential tSH3-binding motif is hard to judge.

To sum up the evaluation of the detected motif hits within known partners of the four NCF1 family proteins (**Table 2 and Table S1**): 1) the binding sites of the tSH3s of NCF1 could likely be better defined within two partners, NCF2 and RelA, 2) although only individual SH3 domains of NCF1 (1st SH3) and TKS5 (1st and 5th SH3) were tested and shown to interact with the cytoplasmic tail of ADAM19, it has a promising tSH3-binding motif candidate, and 3) the interaction between TKS4 and SH3BP1 could depend on the detected tSH3-binding motif within SH3KB1, but individual SH3 domains of TKS4 could also mediate it. Some of the remaining hits are rather unlikely, while others are hard to judge based on the currently available information.

### Sequence signatures of SH3 tandemization and cooperative motif binding within the NCF1 family members

The tSH3s of the four family members are connected by a short, acidic linker, with a relatively well-preserved length within the family (**Figure 3**) and also in the evolutionary sense (**Figure S5**). Only TKS5 is known to have isoform-specific variation within the linker; the first two SH3 domains can only tandemize in the isoform with the shorter linker (UniProt AC: Q5TCZ1-3) [21].

Regarding the domain sequences, the GWW signature amino acid triplets of both domains seem crucial for tandemization, as well as for motif binding [19,20]. The middle Trp of this triplet is well-conserved among SH3 domains [10] and is a key residue in the binding of canonical as well as non-canonical peptide ligands by forming the shallow hydrophobic pockets designated as P_−1_ and P_0_ following the notation by Yu *et al*. [13]. The tryptophanes in this position of both domains contact the strongly conserved Pro in the 2nd position of the tandem SH3-binding motif defined above. The Gly residues in the GWW signatures of both domains are important in tandemization, as larger residues in this position would result in a steric clash and prevent tandemization [19,20]. Accordingly, mutations to Ser in these positions are implicated in autosomal recessive chronic granulomatous disease (CGD) [62]. The last Trp in the triplets of both domains forms a hydrogen bond with the residue two positions ahead of the GWW triplet of the other SH3 domain (**Figure 3**), therefore, they also seem strictly required for tandemization. Interestingly, the hydrogen-bonded residues ahead of the GWW triplets are not fully conserved within the family; in both of these positions, NOXO1 contains residues that are not even similar to the ones observed in other family members (Pro versus Glu in the N-SH3 domain, and D versus L in the B-SH3 domain). Groemping *et al*. proposed that only SH3 domains with the GWW triplet signature could likely form the tandem arrangement [19].

Interestingly, many of the domains’ positions that contact the tandem SH3-binding motif show a remarkable sequence variation within the family (**Figure 3**). Still, some conserved positions can be seen and considered as important motif-binding signatures. Firstly, the C-SH3 domain contains a highly conserved double-negative signature. These acidic residues form salt bridges with the strongly conserved Arg in the 4th position of the tSH3-binding motif. Secondly, the hydrophobic patch at the end of the N-SH3 domain has fully and highly conserved positions that are in contact with the proline residue in the 3rd position of the motif, as well as the residue in the 5th, [RP] position (**Figure 3**). These will be referred to as C-double negative (C-[DE][DE]) and N-PxxΦL (where Φ is an aromatic residue) signatures from now on. It is important to note that several residue positions of these three signatures show remarkable sequence variation among SH3 domains (see SH3 alignment in Table S1 of Teyra J. *et al. [10]*).

The 3rd position of the core motif was defined as a strongly required Pro in the strong motif definition, but it is the one that got somewhat relaxed in the weak motif definition due to sequence variations seen in the weak-binding NOXO1 C-terminal autoregulatory motif. Interestingly, two of the three positions that consistently contact this P(3) residue in the structures show remarkable sequence variation within the NCF1 family (see P(3) contacting several residues in the N-PxxΦL signature in **Figure 3**). This could imply that certain family members are more suited to accommodate motifs with a variation in the 3rd position, while others are more conservative.

The sequence signatures for SH3 tandemization and composite peptide binding introduced above are well conserved for the NCF1 family members among vertebrates (**Figure S5**), along with the respective autoregulatory motifs and partner motifs (**Figure S1**). However, we identified an interesting case of co-evolution between inter-dependent protein interaction modules in the African elephant (Loxodonta africana). Here, the GWW signature triplets of both SH3 domains of NCF1 are disrupted/missing, as well as the autoregulatory (AIR) motif at the C-terminal part of the protein that is responsible for the inactive, closed conformation. Furthermore, the respective motif is also missing from the partner, the NADPH oxidase core complex subunit CYBA, so all interacting modules of the NADPH oxidase regulatory system are affected (**Figure 4**). These findings, along with the complete lack of the NOXO1 gene, strongly suggest that the regulation of the NADPH oxidase is completely altered in the African elephant.

**Figure 4:**
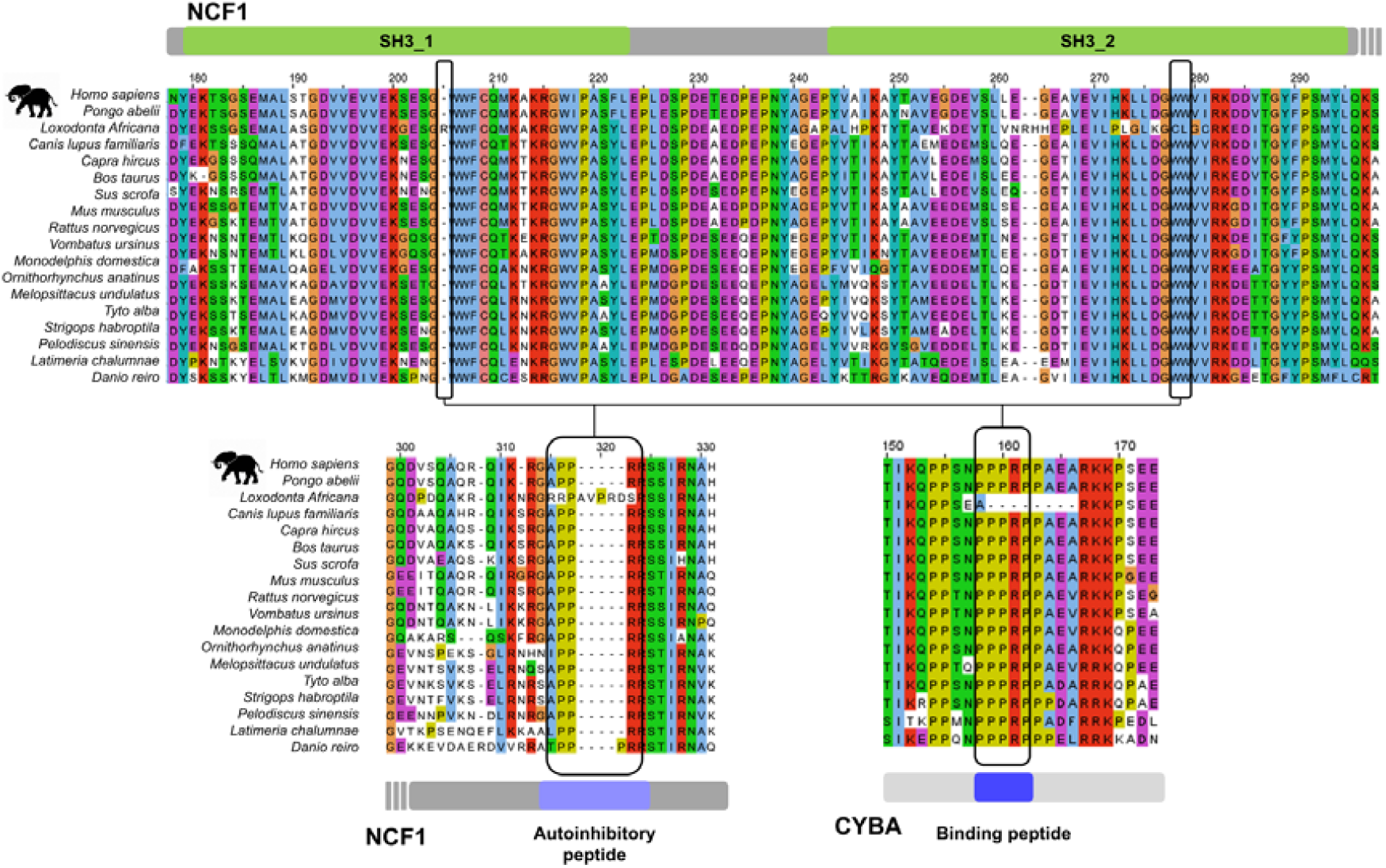
Example for co-evolution between the NCF1 tandem SH3 domains and the respective binding motifs within NCF1 and the partner CYBA in the African elephant (*Loxodonta africana*). Sequence alignments of the NCF1 tandem SH3 domains, the autoregulatory motif and the tSH3-binding motif in CYBA are shown to highlight that all of them got lost or considerably changed in the African elephant (*Loxodonta africana*).

### Does SH3 tandemization and cooperative motif binding occur outside the NCF1 family?

To explore potential candidates for SH3 domain tandemization and cooperative motif binding, we screened all proteins of the human proteome that had at least two SH3 domains (45 proteins; **Table S2**). Interestingly, besides the NCF1 family members and the two close paralogs of NCF1 (NCF1B and NCF1C; could be pseudogenes), only the adaptor proteins SH3KBP1/CIN85 and CD2AP/CMS had at least two consecutive SH3 domains with the GWW signature. While in CIN85 all three domains have the signature, and the first two are relatively close to each other (separated by a linker of ~30 residues [63]), in CD2AP only the last two SH3s have the signature, but those are relatively far apart (~100 residues).

Interestingly, the first two SH3 domains of CIN85 also exhibit the C-[DE][DE] signature and a slightly modified N-PxxΦL signature, with Val in the last position. Therefore, based on SH3 domain signatures, CIN85 could be an interesting candidate for intrachain SH3 tandemization (**Figure S6**). It is important to note, however, that the linker connecting the two domains is much longer than those of the NCF1 family members, and it has a strong basic character at both termini, in contrast to NCF1 family members, where the linker is acidic. This could be a reason why no contacts were identified between the SH3 domains within the CIN85 SH3AB construct (containing the first two SH3 domains of CIN85 connected by the linker) by NMR previously [63]. It is also important to note that there are no isoforms for CIN85 and CD2AP, where the SH3 domains are preserved and only the linker is varied (as for TKS5).

Individual SH3 domains of CIN85 have a non-canonical binding preference for Px[PAV]xPR motifs (where x is any residue) [16] that differs from the binding preference of individual SH3 domains in the NCF1 family [63]. Therefore, even if we assume that they can form tandems and cooperatively bind to peptides, the binding preference might not be the same as for NCF1 family members. We applied different computational approaches to investigate whether the CIN85 SH3 domains could form tandems. First, we used both AF2 and AF3 to predict if the linked first two SH3 domains of CIN85 could bind to a peptide within CIN85 that resembled the tandem SH3-binding motif. CIN85 has a “LPPRR” sequence (residues 417-421 in CIN85; UniProt AC: Q96B97) that though does not fit the strong motif definition due to having a Glu in the C-terminal flanking region (in the 7th position), but might be a weak and/or reverse autoregulatory site. When testing the linked domains (residues 1-160 in CIN85; UniProt AC: Q96B97) with a peptide containing this motif and four flanking residues on both sides, the SH3 domains did not form a tandem arrangement in any of the resulting models (the GWW signatures were too distant to make contacts). Second, we also run AF2 and AF3 with a peptide carrying the proposed autoregulatory motif of the TKS4 protein, because 1) it fits the strong motif definition, 2) the two proteins are binding partners [47,64] and 3) the peptide “QRPVVPPRRPPPP” (residues 753-765 in TKS4; UniProt AC: A1X283) also contains an overlapping PxVxPR motif that fits the binding preference of CIN85 SH3 domains [16] and was recently proved to be the binding site for the close-relative CD2AP [56], implying that it is also the binding site for CIN85. The linked SH3 domains did not form a tandem arrangement in any of the resulting models with this peptide either.

Furthermore, we generated ensembles of 30 conformers using the BioEmu AI-assisted ensemble generation tool [65] on the linked SH3 domains of the NCF1 family members and CIN85 with and without peptides. Without the peptide being part of the constructs, linked SH3 domains failed to sample the typical domain tandem with multiple contacts. However, if the Pro-rich autoregulatory peptide was present C-terminally to the tSH3 domains, the tSH3 protein segments appeared to assemble in a small to large population of the ensemble, depending on the system. In case of NCF1 (aa. 156-310), the interdomain assembly with the peptide appeared most commonly in the correct peptide-bound state. In the TKS4 system (aa. 152-358) and TKS5 system (aa. 166-412), the tandem SH3 was sampled in 19% and 17% of the ensemble population, although plausible peptide binding pose was only detected in the reverse orientation (6% and 17% of conformers, respectively). The NOXO1 construct only sampled the tSH3 in less than 10%, while the autoregulatory peptide was never placed correctly (neither in forward, nor in reverse orientation). While the generated ensembles of NCF1 family members contained varying population with correct tandem arrangements of the SH3 domains (RMSD < 5Å from reference PDB:7YXW) (**Figure S7**), the one generated for CIN85 did not. For the CIN85 system, the addition of the peptide had no detectable effect on the success (p-value=0.35, t-test on corresponding RMSD distributions), despite the fact that this trend of improved tSH3 recovery held true for all other systems (ranging from NOXO1’s p=0.0138 to NCF1’s p=3.8e-17, **Table S3**).

Based on our computational analysis, the N-terminal SH3 domains of CIN85 do not seem to form tandems. Since, based on sequence signatures and domain distances, CIN85 seemed to be the most promising candidate from the whole proteome, our results imply that intrachain tandemization of SH3 domains might only occur within the NCF1 family.

While probably not forming tandems intramolecularly, the N-terminal SH3 domain of CIN85 has already been described to undergo intermolecular clustering with a proline-arginine-rich peptide of CBL-b (PDB: 2BZ8) [66,67]. In these clusters, two different CIN85 proteins are pulled together by motif-binding-mediated clustering of their N-terminal SH3 domains. After identifying this structure and some other similar structures based on Desrochers G *et al*. [68], we performed a targeted search of the PDB for structures wherein there are at least two SH3 domains binding to the same peptide. Only eight structures fulfilled this criterion, including the five depicting the tandem SH3 domains of NCF1 binding to the autoinhibitory or CYBA peptides **(Table 1)**. In PDB ID:2BZ8 [66], introduced above, the two copies of the N-terminal SH3 domain of CIN85 bind to a single CBL-b peptide. In PDB:2D1X [69], two copies of the C-terminal SH3 domain of cortactin bind to a peptide of AMAP1. Finally, in PDB:5SXP [68], two copies of the ARHGEF7/beta-PIX SH3 domain bind to a peptide of the E3 ubiquitin ligase ITCH. Overall, the latter two novel cases also represent interchain SH3 clusterization events, as seen for the CIN85 N-terminal SH3 domain.

When comparing structures depicting intrachain tandemization and interchain clusterization (**Figure 5**), we found some important differences. In the case of tandemization (structures are only available for NCF1), the two domains are in contact and form a common binding groove (buried surface area between the two domains is 572 Å^2^ [20]), where the GWW signature triplets and surrounding residues of both domains play a key role [19,20] (**Figure 3**). In contrast, in the structures depicting interchain SH3 clustering, the two SH3 domains do not form an interface, although they do have the GWW signature, their GWW triplets are much further away, and they bind the peptide independently, i.e. through two independent binding grooves (**Figure 5**).

**Figure 5:**
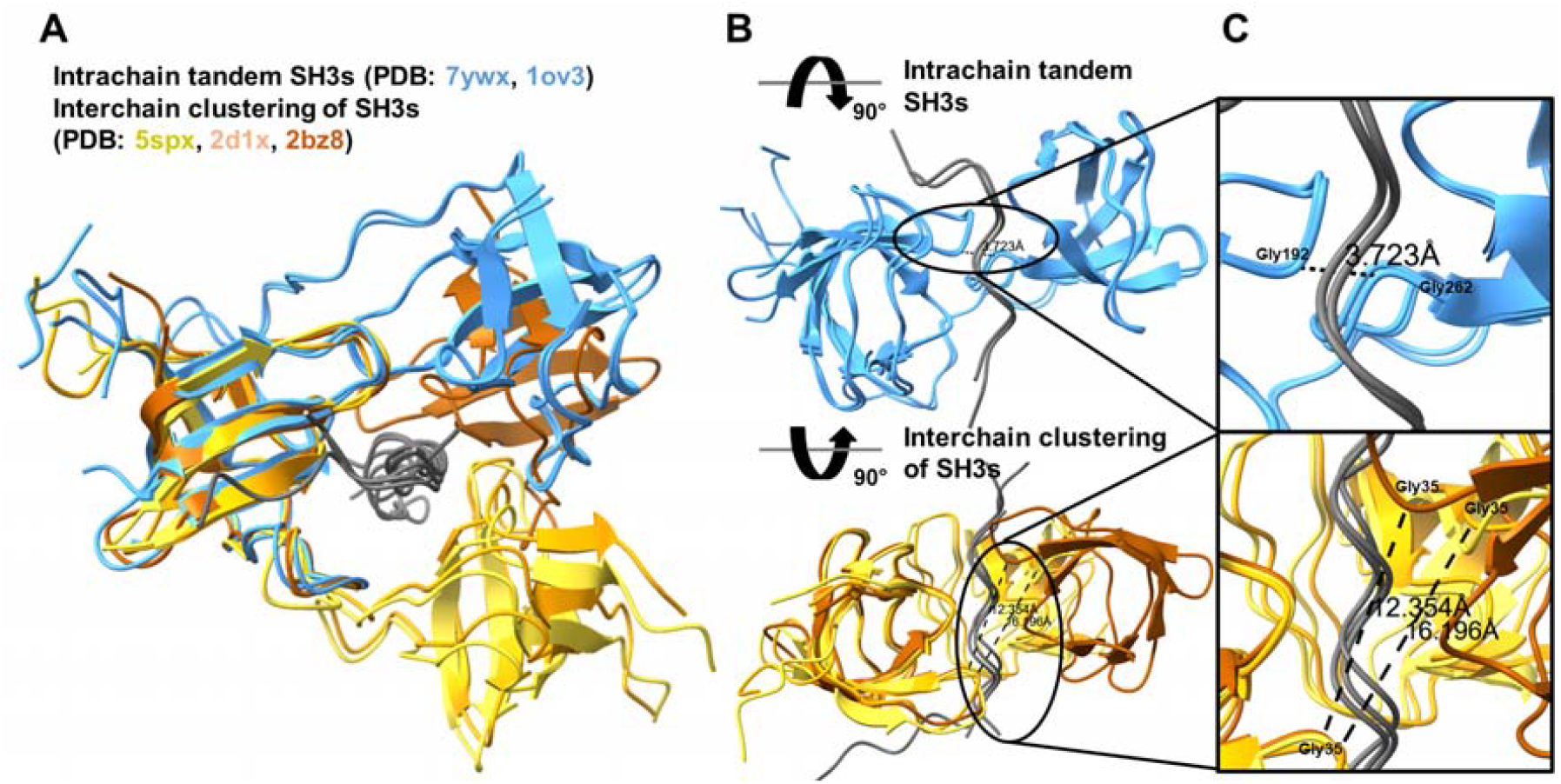
Structure alignment of intrachain tandem SH3 domains and interchain SH3 clustering. A) structure alignment of intrachain and interchain experimental structures (7ywx, 1ov3, 5spx, 2d1x, 2bz8). B) Rotated view on intrachain tandem (top) and interchain clustering (bottom) SH3 domains. C) A closer view highlighting the distance of SH3 domains in intrachain tandem (top) and interchain clustering (bottom) SH3 domains. The depicted distances are calculated between the Gly residues of the GWW signature triplets of the two domains.

While our results suggest that CIN85 does not form intramolecular SH3 domain tandems, and thus those are likely specific to the NCF1 family, they underscore the diversity of possible interactions between SH3 domains. The observed interchain clustering of the CIN85 N-SH3 domain indicates that SH3 domain interactions are context-dependent, though the full scope, associated sequence signatures and mechanisms of these effects remain to be fully understood.

## Discussion

SH3 domains are small but largely adaptable protein modules that evolved to fulfil a versatility of molecular functions [2]. The ~300 SH3 domains of the human proteome [1] display a fascinating variety of binding preferences [1,9,10], and thereby bring specificity and fidelity into cellular signaling. While most known SH3 domains follow a simple one-to-one peptide binding mode, there are exceptions to this rule. For instance, the IB1 SH3 domain is not known to bind any peptides, the residues usually involved in peptide binding instead form a unique dimerization interface [70]. In our study, we focused on a unique binding mode wherein two tandemly arranged SH3 domains of a single protein chain come together to form a common binding groove that accommodates a single Pro-rich peptide [18–20]. This binding mode has only been described for the NCF1 family of proteins, NCF1 [18–20,23,24,71,72], NOXO1 [25,30,31], TKS4 [26] and TKS5 [21,27]. While for NCF1 it has already been discovered and characterized in much detail decades ago [18–20,23,24,71,72], for the last family member, TKS4, it has only been verified recently [26]. Through the years more and more evidence and examples accumulated, but – probably due to the big differences in timing and the imbalance of available data for the different family members – those remained scattered in the literature and have never been collected and comprehensively analyzed.

Here, we collected the scattered pieces of evidence, which enabled taking a fresh look at the gathered data, systematic cross-comparison and contextualization of published results across proteins and deriving novel insights. Definition of the tandem SH3-binding motif as a novel short linear motif (SLiM) based on the gathered instances was a crucial step towards identifying novel motif candidates. We relied on the classical approach for the definition of the tSH3-binding motif [28,29] and detection of potential motif instances in the human proteome. Due to their short length and few specificity-determining residue positions, motif definitions are degenerate and tend to occur in protein sequences by chance [37]. This burdens pattern-based motif searches with high false positive rates [37]. Therefore, besides applying the usual restrictions to get rid of false positives, such as only considering motifs within disordered regions [28,39], we also narrowed down our hits to those occurring in the known binding partners of NCF1 family proteins, and manually curated the resulting list.

We applied state-of-the-art AlphaFold-based structure prediction approaches for assessing the validity and likely binding modes of both, validated motif instances with no available structures and novel motif instances proposed in our study. We applied the AlphaFold2-derived AlphaFold-Multimer method [73], which demonstrated high accuracy in predicting protein complexes. Although folding and binding are based on similar biophysical principles, and the method was primarily expected to excel in predicting domain interactions, it can also reliably predict disordered regions bound to ordered domains and distinguish true interactions from decoys [74]. Due to their short length and preferential localization in IDRs with poor sequence conservation, predicting the complex structures for SLiM-mediated interactions is particularly challenging. They may fall outside AlphaFold’s training set, and it is not clear to what extent can AlphaFold apply co-evolutionary signals for their prediction. Nevertheless, AlphaFold-Multimer has shown success in both systematically screening known interactions [75] and, in some cases, discovering new motif instances [76]. Furthermore, it has been shown to capture subtle differences in the binding modes of linear motifs [77,78], a feature that is particularly important for the evaluation of the versatile peptide binding modes of SH3 domains. A crucial feature for any prediction algorithm is to provide reliability or probability metrics for its predictions. While AlphaFold-Multimer provides predicted TM-scores (ptm), these scores are often biased towards the much larger globular domains, while failing to represent short motifs. To overcome this issue, we also considered actifpTM [79] confidence scores that are specifically adapted to measure the reliability of domain–peptide interactions. Although AlphaFold3 [80] is expected to perform well on motifs, no large-scale analysis has demonstrated this so far, therefore we only used it as a complementary approach.

We also experimented with the recently developed BioEmu deep learning-based ensemble generation tool [65] to obtain conformational ensembles for the tandem SH3 domains of the NCF1 family members with, or without the respective autoregulatory motifs linked to the tandems. The rationale for using BioEmu was that it not only generates AlphaFold-like structural models based on a training set of rigid protein structures, but is also trained on extensive molecular dynamics (MD) simulations and complementary experimental stability data from small proteins [81]. This combination makes BioEmu particularly valuable for investigating kinetic observables, such as the conformational dynamics of domain-domain interactions and peptide-induced conformational changes. Interestingly, we observed that BioEmu often responds sensitively to the addition of autoregulatory Pro-rich peptides, which appear to promote tandemization of SH3 domains in NCF1 family proteins. It remains unclear whether tSH3 formation facilitates peptide binding, or whether peptide binding itself induces tandemization of the SH3 domains in the cellular context. This question has not been thoroughly investigated, and targeted kinetic studies will be necessary to elucidate the mechanisms underlying this intramolecular assembly.

Our structure prediction approach was reinforced by the observation that the complexes of tSH3s and motifs fitting our strong motif definition were predicted with almost identical structures as the experimentally determined complexes. At the same time, it highlighted the possibility that certain tSH3-binding motifs, seemingly the ones with at least one negatively charged residue in the two C-terminal flanking positions immediately following the core motif, bind in a reverse orientation. We suggest that this novel binding mode observed for the N-terminal autoregulatory tSH3-binding motif of TKS4 [26] and the homologous motif proposed for TKS5 autoregulation should be experimentally investigated.

When screening the proteome for candidates of the tSH3 binding mode using the previously established sequence signatures of tandemization and cooperative motif binding, SH3KBP1/CIN85 was identified as the only promising candidate. However, in accordance with the results of a previous structural analysis by NMR [67], our structure prediction and ensemble generation approaches did not support the formation of the tandem arrangement, even in the presence of Pro-rich peptides harboring the tSH3-binding motif, or a PxxxPR motif fitting the binding preference of individual CIN85 SH3 domains. Based on these observations we conclude that the tandem SH3-binding mode is likely to be specific for the four members of the NCF1 protein family that have two or more SH3 domains.

We successfully identified promising candidate tSH3-binding motifs in both NCF1 family members (representing potential autoregulatory sites) and in their known binding partners. Since most NCF1 family members are directly implicated in diseases, the proposed novel autoregulatory mechanisms and mapped binding sites within partner proteins could have therapeutic relevance. Mutations preventing the tandemization of SH3 domains within NCF1 [19,20,62] or destroying the tSH3-binding motif in the partner CYBA [18] lead to a lack of functional NADPH oxidase and are implicated in Chronic Granulomatous Disease (CGD), a severe immune deficiency. Our proposed motifs (**Tables 1 and 2**) include a potential secondary autoregulatory site within NCF1 (overlapping with the C-terminal Pro-rich binding site of the NCF2 SH3 domain [48]) and a potential tSH3-binding motif within NCF2 that could represent a secondary binding interface between these two crucial subunits of the NADPH oxidase [18,48]. Furthermore, we believe that with the help of our motif definition, we managed to precisely map the NCF1 tSH3-binding site within the protein RelA, which mediates the NCF1-RELA interaction underlying the RelA-mediated activation of the NF-κB pathway by IL-1 in endothelial cells [49].

The role of the tSH3 domains within the larger family members, TKS4 and TKS5 is much less understood [21,26,27,82], although both of them are heavily implicated in diseases, most importantly in the metastatic potential of different cancers, such as melanoma, lung cancer and colon cancer, among others [32,33,47,56,82–84]. Having both overlapping and specific functions, the two proteins are key in the formation and dynamic changes of podosomes and invadopodia [34,82]. They are important for the degradation of the extracellular matrix around these cell protrusions (through facilitating the transport of matrix metalloproteases to the cell surface [82] and directly communicating with them in the cytoplasm [50,51]), which helps free up the space required for cell migration. At the same time, they also play a role in the remodeling of the cytoskeleton inside the protrusions, e.g. TKS4 can directly interact with the actin capping protein [47] (and also indirectly through its interaction with CD2AP [56,85]), and therefore likely plays a key role in reorganizing the actin cytoskeleton at podosomes and invadopodia. Due to the crucial roles of TKS4 and TKS5 in the metastatic potential of cancer cells [82], understanding the regulatory mechanisms affecting their availability and/or functions, as well as the molecular mechanisms of their protein-protein interactions, could have direct therapeutic relevance.

We propose several novel candidate tSH3-binding motifs that could be important in the regulation and functioning of the two proteins (**Tables 1 and 2**). First, we propose an autoregulatory tSH3-binding motif between the 3rd and 4th SH3 domains of TKS4 that was previously uninvestigated, despite its sequence and AF2 structure prediction indicating it to be a stronger site than the one previously suggested [26]. A further interesting detail is that the recently described CD2AP-interacting SH3-binding motif, PxVxPR [56] completely overlaps with this proposed downstream autoregulatory site of TKS4. This suggests that if the proposed intramolecular interaction exists, it could likely be resolved by the binding of CD2AP. However, the validity and biological relevance of this potential autoregulatory mechanism still awaits experimental validation.

Our structure predictions and ensemble calculations also suggest that the previously suggested autoregulatory tSH3-binding motif falling between the 2nd and 3rd SH3 domains of TKS4 [26] likely binds in reverse orientation (just as the homologous site within TKS5 that could be an autoregulatory motif of the short isoform), probably due to having a negatively charged residue in the first position of the C-terminal flank. While candidate tSH3-binding motifs have also been detected in a partner of TKS4 (SH3BP1 [47]) and in a partner of TKS5 (ADAM19 [50]), these motifs lie in long Pro-rich IDRs that harbour many potential SH3-binding sites, thus the interactions could also be mediated by the individual SH3s of the two proteins. In the case of ADAM19, the 1st and 5th SH3 domains of TKS5 were demonstrated to mediate the interaction [50]. Nonetheless, both the potential autoregulatory and partner binding candidate tSH3-binding motifs are proposed for experimental validation, wherein, ideally, constructs for both, tandem and individual SH3 domains should be used in order to fully elucidate the participating protein modules and the underlying specificity determinants of the interactions.

## Conclusion

In our study, we revisited a unique binding mode described for the p47^phox^-related protein family, wherein two tandemly arranged SH3 domains of a single protein chain come together to form a common binding groove that accommodates Pro-rich peptides. When screening the human proteome for other potential candidates of SH3 tandemization relying on the associated specific sequence signatures, only one promising candidate could be identified. However, our structure prediction and ensemble generation approaches did not support the formation of the tandem arrangement by the SH3 domains of the one candidate, so we conclude that this binding mode is likely specific to the NCF1 protein family. Through the collection and comprehensive analysis of previously described tandem SH3-binding motif instances scattered in the literature, we successfully defined the binding preference of the tandem SH3 domains as a novel short linear motif (SLiM), [PAVIL]PPR[PR][^DE][^DE]. This motif definition was then used to discover novel motif instances within NCF1 family members and their interaction partners. The resulting hits were manually curated and their validity and potential binding modes evaluated by state-of-the-art AI-assisted structure prediction and ensemble generation approaches. Our results imply that some specific instances of the tSH3-binding motif bind in a novel, reverse binding mode, and propose this binding mode along with the most promising candidate motif instances discovered for experimental validation. Due to the involvement of several members of the studied protein family in different diseases, especially the TKS4 and TKS5 proteins being heavily implicated in the metastatic potential of diverse cancer types, the proposed potential autoregulatory and partner-binding tandem SH3-binding motifs could have direct therapeutic relevance.

## Materials and Methods

### Identification of tandem SH3 domains

To identify SH3 domain-containing proteins, we searched for human proteins with the PF07653, PF00018, PF14604 PFAM domain identifiers in the InterPro database [86]. Domain boundaries (taken from InterPro annotations) were evaluated and selected as tandems if the linker region connecting two SH3 domains was shorter than 60 amino acids. In the case of the NOXO1 protein (a member of the NCF1 family that is in the focus of this work), one of the two SH3 domains was missing from the InterPro annotations, and therefore it needed to be added based on UniProt annotation (where it was present in accordance with the literature).

### Multiple sequence alignments

Orthologous proteins were collected from OMA [87] (using Uniprot AC identifiers) and selected to uniformly cover Vertebrates (**Table S4)**. Sequences were aligned with ClustalΩ [88] and visualized in Jalview [89].

### Search for structures and structural analysis

Possible tandem structures were collected using different methods. The three PFAM IDs belonging to SH3 domains (PF07653, PF00018, PF14604) were searched in 3did [90] to retrieve SH3-containing entries. Each entry was manually assessed to filter those where multiple SH3 domains occur and a peptide is part of the complex. To validate the results using an independent method, the tandem structures found were also used as an input in FoldSeek [91], and other tandem SH3s were searched in the PDB100 dataset [92]. From the available structures, the domain-peptide amino acid level interactions were predicted using RING [93] with default settings.

### Structure prediction

We used ColabFold 1.5.5 [94] (based on AlphaFold2 (AF) [42]) to predict the tandem domain-motif interactions defined in **Table 1**. We opted AF2 over AF3 for the following reasons: 1) several high-throughput analyses concluded that AF2 can reliably predict motif-domain interactions [75,76], however such an investigation is not available for AF3; 2) In ColabFold we can directly manipulate which template structure is used by the prediction; 3) A modification for AF2 can produce actifpTM [79], a score directly developed to assess the quality of motif-domain interactions; 4) We have previously shown that this methodology can capture slight deviations in binding mode [78]. For each interaction, we used PDB:7yxw as a template and added the sequence of the tandem SH3 domain together with the core motif (+-4 positions). Among the predicted models, we selected the one with the highest actifpTM (**Table S5**). We used ChimeraX [95] for visualization. In edge cases (reverse motif orientation, SH3KBP1/CIN85 clustering), we also confirmed the structure prediction using AF3 [80].

### Conformational ensemble prediction

To emulate the structural ensemble of the SH3 tandems with and without the Pro-rich binding motifs, the ColabFold version [96] of BioEmu 1.1 [65] was used with the following parameter setup: Number of samples: 30. Number of samples to randomly select for clustering: all available samples. The coverage threshold used for FoldSeek [91] clustering was set to 0.7, with a minimum TM-score of 0.6, and 95% sequence identity threshold used for clustering. Shorter constructs only constituting the two SH3 domains were tested using the following region borders (UniProt numbering): for NCF1: aa. 156-285, for NOXO1: aa. 163-296, for SH3KBP1/CIN85: aa. 1-157, for TKS4: aa. 152-280 and for TKS5 isoform 3: aa. 166-297. Longer constructs also comprising the Pro-rich peptide were the following: NCF1: aa. 156-310, NOXO1: aa. 163-343, SH3KBP1/CIN85: aa. 1-157 fused to aa. 334-432, TKS4: aa. 152-358 and TKS5 isoform 3: aa. 166-412. For the comparison of conformers to tandem SH3 reference crystal structure PDB:7YXW, PyMOL’s (ver. 3.1.1., Schrödinger LLC) superimposition tool was used: ‘super sele, topology.pdb, cycles=0, target_state=i’, where sele corresponds to 7YXW’s PDB region 160-211 and 229-283, and ‘i’ is a given conformer of topology.pdb.

### Motif prediction

We scanned the canonical human proteome (obtained from UniProt [43], 2025_1 release) for the defined strong binding motif ([PAVIL]PPR[PR][^DE][^DE]). We calculated or added the following properties: 1) protein disorder [46]. 2) We defined motif conservation using the regular expression. We used a set of Chordata species proteomes (**Table S6**) and searched orthologs for human proteins using BLAST (the sequence with the highest sequence identity (above 20%) was selected for each species, if it covered at least 60% of the human sequence), and then aligned the selected sequences with ClustalΩ [88]. We searched for the regular expression in the orthologous proteins. Due to the challenging alignment of disordered regions, we allowed 50-residues deviation in the alignment for regular expression hits. 3) Broad-level localization definition of the motif containing protein (i.e., defining intracellular proteins using the same methodology as was used during the development of TOPDOM [97]). 4) To identify if the motif hit is likely to occur simply due to high Pro/Arg content of the sequence, we measured how many times the motif is found after randomly shuffling the protein sequence 100 times. 5) Whether the identified protein is listed as a partner of NCF1, NOXO1, TKS4 or TKS5 in BioGRID [98] and/or IntAct [45]. Additional partners of the TKS4 protein were derived from a recently published dataset [47].

## Supplementary Materials

### Supplementary Figures are available in the Supplementary Material

**Figure S1: Sequence alignments of the experimentally verified and proposed motifs from Table 1 on vertebrate species**. Only the proximity of the motifs are depicted and the motifs themselves are boxed.

**Figure S2: Structure alignment showing the correspondence between experimental and AF2-predicted structures**. The experimental structures (1ov3, 1wlp, 7yxw) of tandem SH3 domains with the SH3-binding motifs are depicted in blue, while the AlphaFold2-predicted structures of motifs fitting the strong motif definition (strong binding) are shown in green.

**Figure S3: AlphaFold3 models of the tandem SH3 domains with their autoinhibitory peptide(s)**. The NCF1 1st and 2nd, as well as the TKS4 2nd motifs that fit the strong motif definition bind in forward orientation, while the TKS4 1st and TKS5 motifs bind in reverse orientation (the directions correspond to the ones predicted by AF2). The weak-binding NOXO1 motif is not placed into the same binding groove as the other motifs. The experimentally validated autoregulatory motif instances are marked by a star, the others are proposed instances.

**Figure S4: Sequence alignments of the proposed motifs within binding partners from Table 2 on vertebrate species**. Only the proximity of the motifs are depicted and the motifs themselves are boxed.

**Figure S5: Sequence alignments of the tandem SH3 domains of NCF1 family members**. Domain boundaries of SH3 domains are highlighted.

**Figure S6: Sequence alignment of the tandem SH3 domains in NCF1 family members and CIN85**. The alignment shows that while the GWW, C-double negative (C-[DE][DE]) and N-PxxΦL tandemization and tSH3 motif binding signatures are well-conserved in CIN85, the linker is much longer and oppositely charged compared to NFC1 family members.

**Figure S7: Structural superposition statistics of the BioEmu predicted ensembles with the reference state of tandem SH3 domain**. Structural superpositions of (A) NCF1, (B) NOXO1, (C) TKS4 and (D) TKS5–3 ensemble conformers with the chosen reference structure of NCF1 tandem SH3 solved by X-ray crystallography at resolution of 2.5Å.

### Supplementary Tables

**Table S1:** List of strong tandem SH3-binding motif instances in the human proteome. Protein disorder, conservation, localization, occurrence of motifs by chance in the protein sequence and NCF1 family member interactions are also listed.

**Table S2:** List of SH3 domains (with and without GWW signature), and their distance from each other.

**Table S3**: Structural superposition results and corresponding statistics of the BioEmu predicted ensembles with the reference state of tandem SH3 domain.

**Table S4:** List of species investigated on the alignment figures.

**Table S5:** ActifpTM scores of AF2-predicted structures from Table 1.

**Table S6:** List of species considered for calculation of sequence conservation for the strong tandem SH3-binding motifs detected in the human proteome.

## Author Contributions

Conceptualization, R.P., Z.E.K. and L.D.; Methodology, R.P., Z.E.K., L.D. and T.L.; Formal Analysis, Z.E.K., L.D., T.L. and R.P.; Investigation, Z.E.K., R.P. and L.D.; Data Curation, R.P. and Z.E.K.; Writing – Original Draft Preparation, R.P., L.D., Z.E.K. and T.L.; Writing – Review & Editing, R.P., Z.E.K., L.D. and T.L.; Visualization, Z.E.K., L.D., T.L. and R.P.; Funding Acquisition, R.P., L.D. and Z.E.K.

## Institutional Review Board Statement

Not applicable.

## Informed Consent Statement

Not applicable.

## Data Availability Statement

The original contributions presented in this study are included in the article/supplementary material. Further inquiries can be directed to the corresponding author.

## Abbreviations

AF: AlphaFold
AIR: Autoinhibitory region
CGD: Chronic granulomatous disease
ELM: Eukaryotic linear motif resource
IDR: Intrinsically disordered region
ITC: Isothermal titration calorimetry
PDB: Protein data bank
PPI: Protein-protein interaction
PPII: Polyproline type II
SLiM: Short linear motif
tSH3s: tandem
SH3: domains

