## Supplementary material for "Definition and discovery of tandem SH3-binding motifs interacting with members of the p47^phox^-related protein family": Figure S

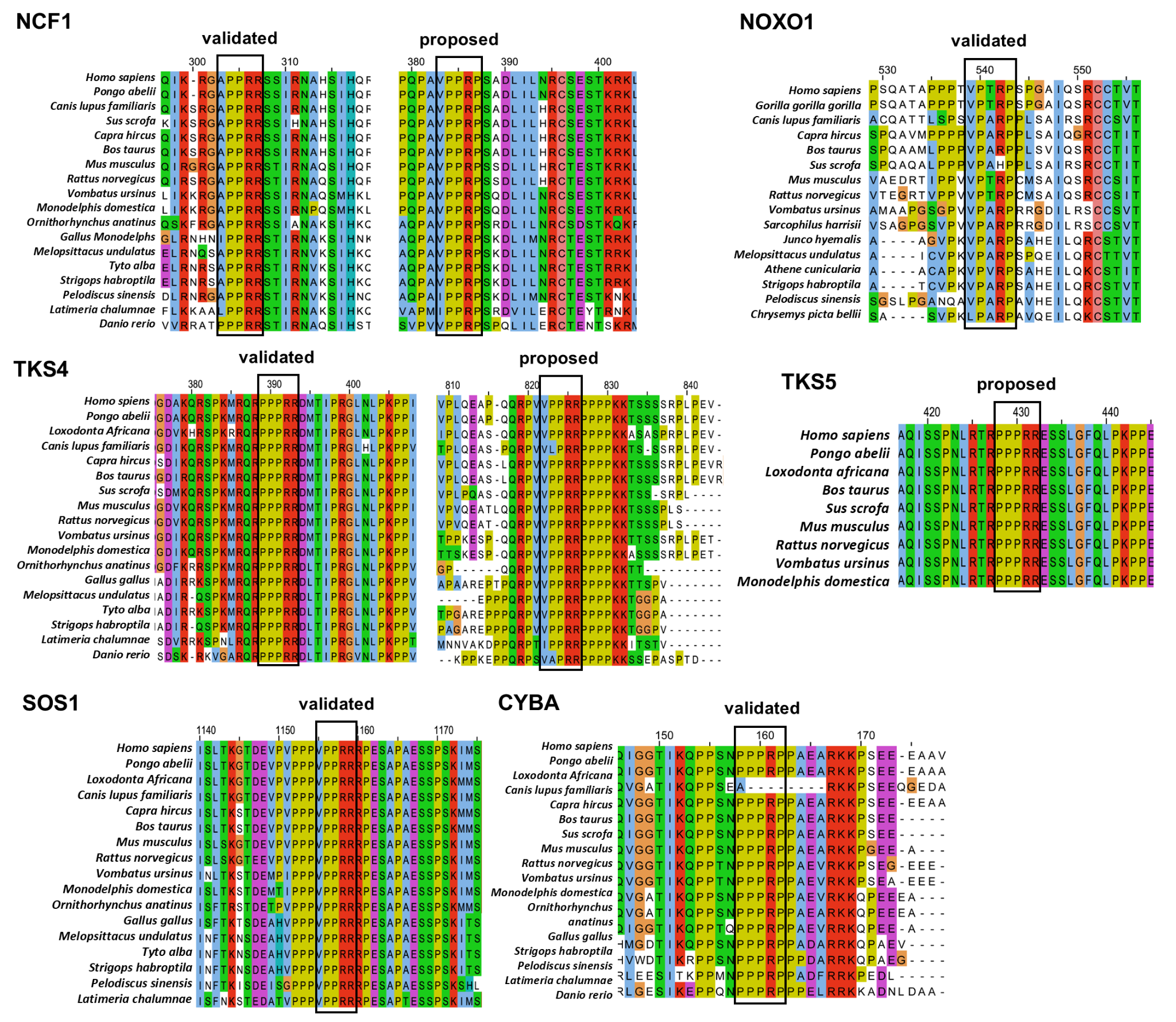

**Figure S1: Sequence alignments of the experimentally verified and proposed motifs from Table 1 on vertebrate species.** Only the proximity of the motifs are depicted and the motifs themselves are boxed.

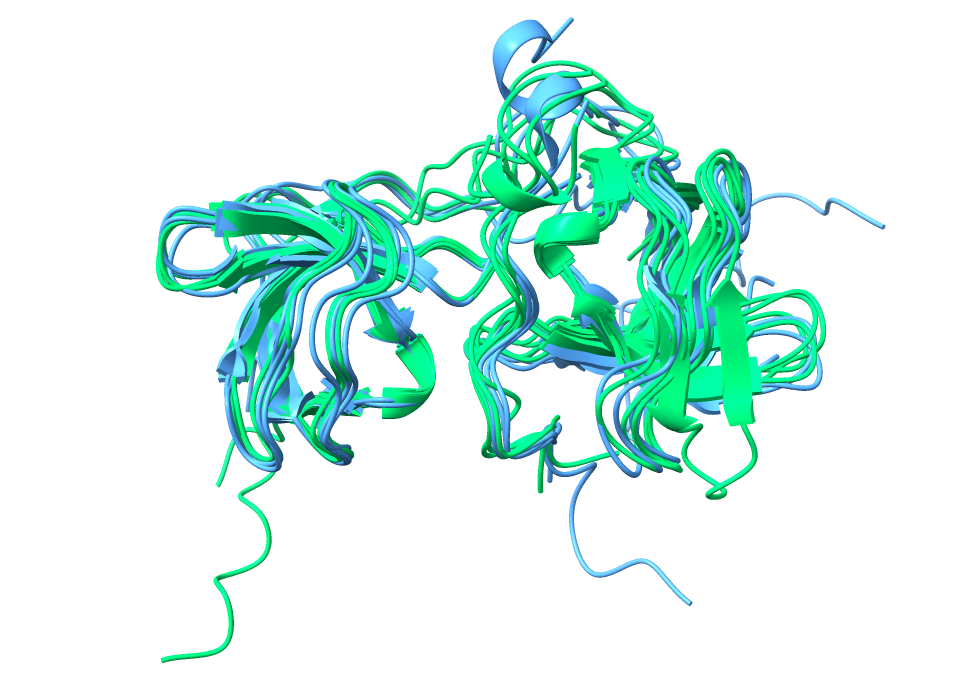

**Figure S2: Structure alignment showing the correspondence between experimental and AF2-predicted structures.** The experimental structures (1ov3, 1wlp, 7yxw) of tandem SH3 domains with the SH3-binding motifs are depicted in blue, while the AlphaFold2-predicted structures of motifs fitting the strong motif definition are shown in green.

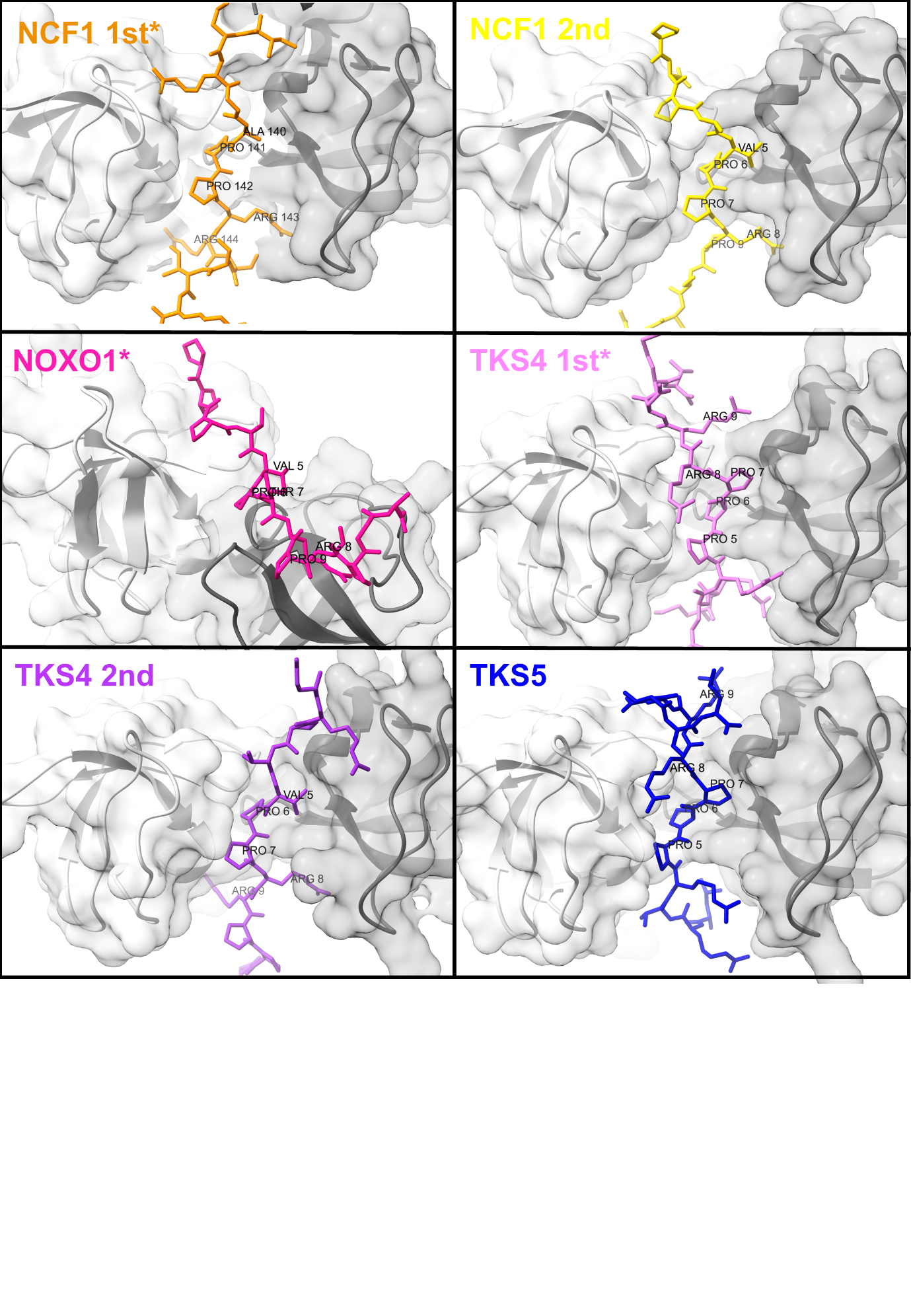

**Figure S3: AlphaFold3 models of the tandem SH3 domains with their autoinhibitory peptide(s).** The NCF1 1st and 2nd, as well as the TKS4 2nd motifs that fit the strong motif definition bind in forward orientation, while the TKS4 1st and TKS5 motifs bind in reverse orientation (the directions correspond to the ones predicted by AF2). The weak-binding NOXO1 motif is not placed into the same binding groove as the other motifs. The experimentally validated autoregulatory motif instances are marked by a star, the others are proposed instances.

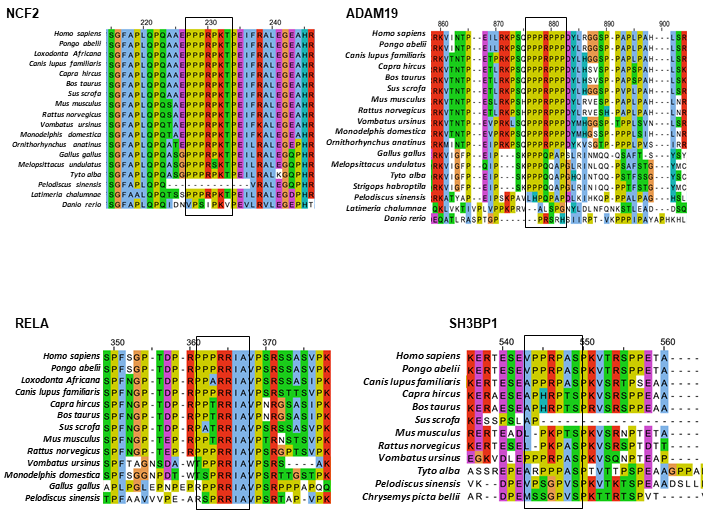

**Figure S4: Sequence alignments of the proposed motifs within binding partners from Table 2 on vertebrate species.** Only the proximity of the motifs are depicted and the motifs themselves are boxed.

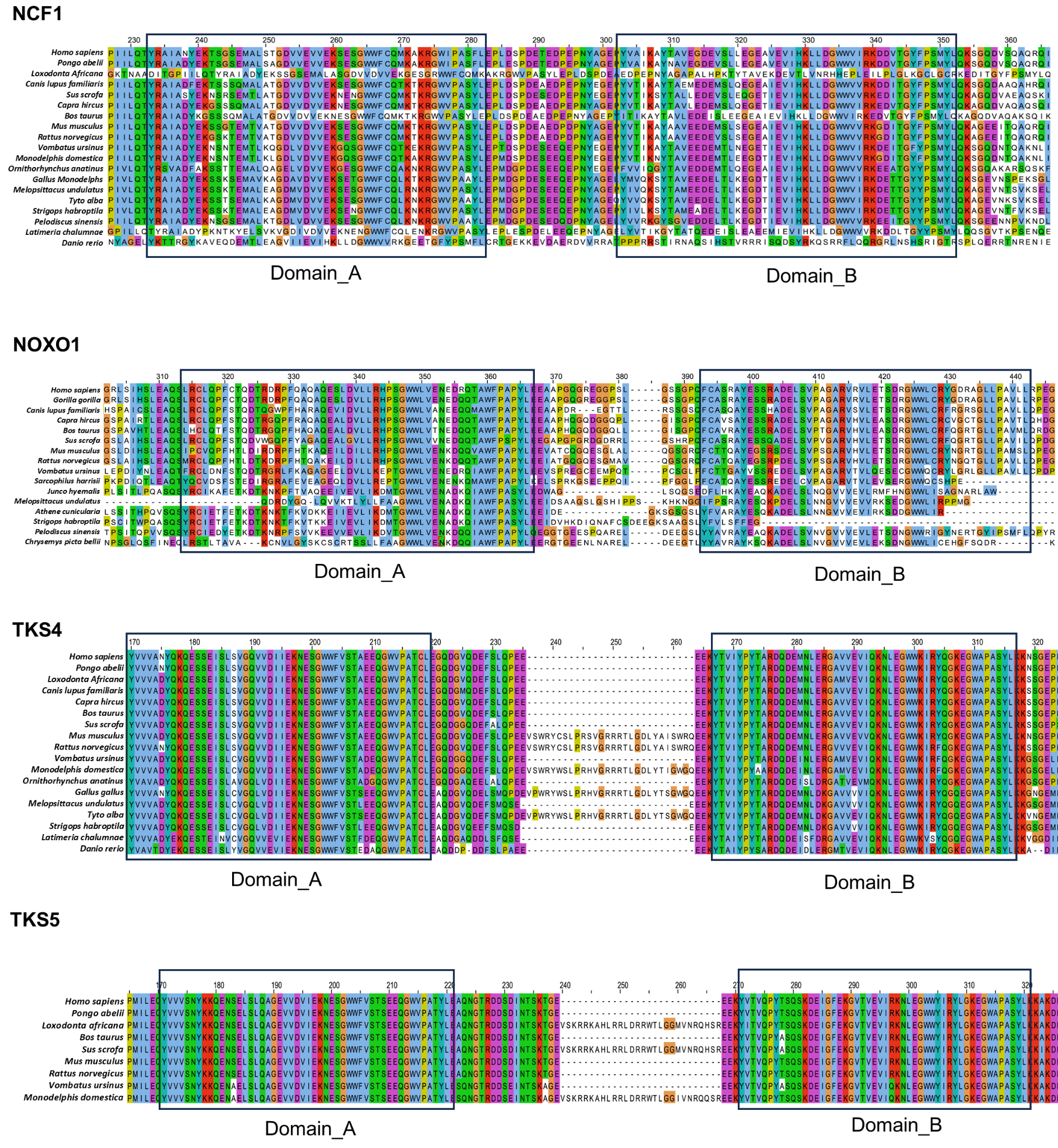

**Figure S5: Sequence alignments of the tandem SH3 domains of NCF1 family members.** Domain boundaries of SH3 domains are highlighted.

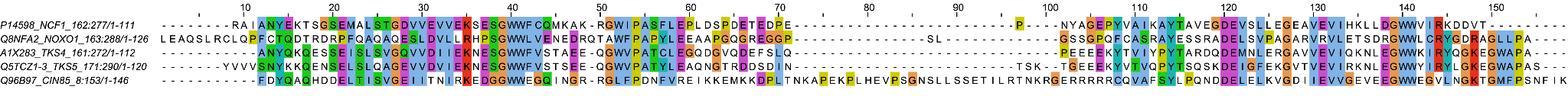

**Figure S6: Sequence alignment of the tandem SH3 domains in NCF1 family members and CIN85.** The alignment shows that while the GWW, C-double negative (C-[DE][DE]) and N-PxxΦL tandemization and tSH3 motif binding signatures are well-conserved in CIN85, the linker is much longer and oppositely charged compared to NFC1 family members.

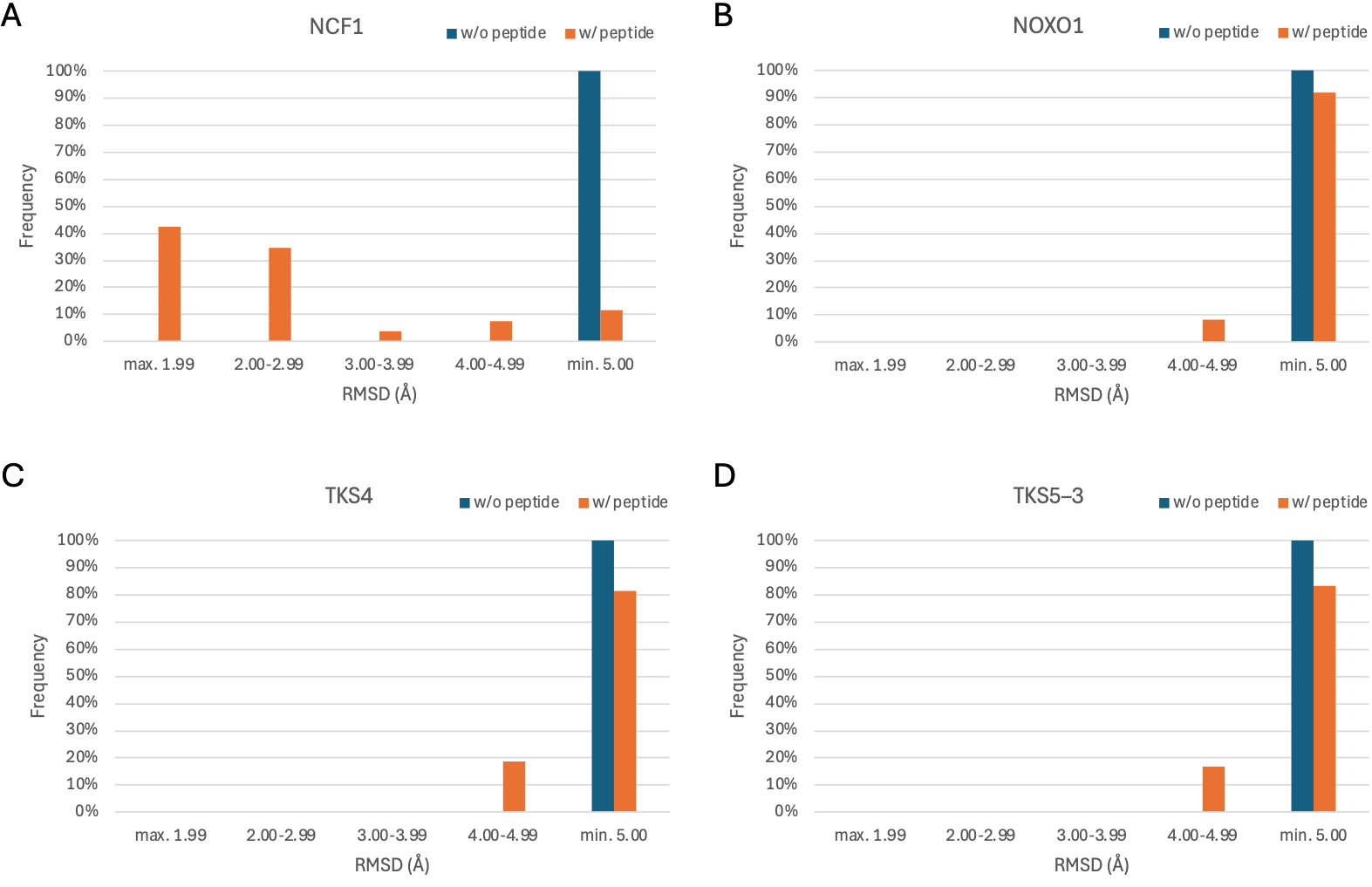

**Figure S7: Structural superposition statistics of the BioEmu predicted ensembles with the reference state of tandem SH3 domain.** Structural superpositions of (A) NCF1, (B) NOXO1, (C) TKS4 and (D) TKS5–3 ensemble conformers with the chosen reference structure of NCF1 tandem SH3 solved by X-ray crystallography at resolution of 2.5Å.
